# Restriction of HIV-1 infectivity by interferon and IFITM3 is counteracted by Nef

**DOI:** 10.1101/2025.05.15.654345

**Authors:** Mahesh Agarwal, Kin Kui Lai, Isaiah Wilt, Saliha Majdoul, Abigail A. Jolley, Mary Lewinski, Alex A. Compton

## Abstract

The viral accessory protein Nef is a major determinant of HIV-1 pathogenicity in vivo. Nef is a multifunctional, immunomodulatory protein that downmodulates cell surface proteins, including CD4 and MHC class I (MHC-I) important for T-cell-mediated immunity. In addition, Nef also regulates cell-intrinsic immunity—Nef boosts the infectivity of virions produced and released from HIV-infected cells, at least in part, by counteracting the antiviral activity of transmembrane proteins SERINC3 and SERINC5. However, it has been reported that the enhancement of HIV-1 infectivity by Nef persists in certain cell lines deficient for SERINC3/5, revealing the existence of other Nef-sensitive host factors that impact HIV-1 infectivity. Here, we show that Nef proteins, especially those derived from many primary isolates of HIV-1, restore infectivity in interferon-treated cells and confer resistance to interferon-induced transmembrane protein 3 (IFITM3). IFITM3 is a restriction factor that reduces retroviral infectivity by incorporating into virions, inhibiting Envelope glycoprotein function, and reducing entry into cells. Using a primary isolate of Nef derived from HIV-1 clade C, we found that Nef interacts with IFITM3 in membranes and uses the endocytic adaptor protein AP-2 to counteract it. Furthermore, Nef reduced IFITM3 cell surface levels, increased IFITM3 levels in early endosomes, and reduced IFITM3 incorporation into HIV-1 virions. Nef also impaired IFITM3 oligomerization and restored membrane fluidity in IFITM3-expressing cells. The antiviral activity of IFITM3 and its inhibition by Nef were unaffected by SERINC5 knockdown, suggesting that the counteraction of IFITM3 represents a unique function of Nef. Our findings reveal a previously unrecognized immunomodulatory role for Nef in the setting of the interferon-induced antiviral state during HIV-1 infection.

## INTRODUCTION

The viral accessory protein Nef is a major determinant of HIV-1 pathogenicity and is found in all members of the primate lentivirus family. Although non-essential for HIV-1 infection and replication in most tissue culture models, in vivo studies in humans and non-human primates showed that Nef contributes to disease progression (1, 2). Nef is a myristoylated protein that associates with cellular membranes and regulates cellular signaling and vesicular trafficking pathways (3). By acting as a molecular bridge that links cellular cargo to the vesicular transport machinery, including the clathrin adaptors (AP-1, AP-2, and AP-3), Nef alters the abundance of a growing list of cell surface proteins (4, 5). These include the immune-related cell surface proteins CD4 and MHC class I, which function in T cell signaling and antigen presentation, respectively (6–8).

Another well-characterized yet incompletely understood function of Nef is its ability to boost the infectivity of virions produced and released from HIV-infected cells. The Nef-mediated enhancement of viral infectivity has been attributed, at least in part, to counteraction of transmembrane serine incorporator (SERINC) family members SERINC3 and SERINC5, whose downmodulation and exclusion from HIV-1 virions is associated with enhanced virus fusion (9, 10). The significance of HIV-1 restriction by SERINC3/5 and its antagonism by Nef is supported by the demonstration that the global spread of HIV-1 correlates with the potency of SERINC5 counteraction (11). Furthermore, Nef from primary isolates display elevated activity against SERINC as compared to Nef from the lab-adapted strain NL4-3 (12), and Nef isolated from HIV-1 controllers was shown to antagonize SERINC5 poorly compared to Nef isolated from HIV-1 progressors (13). Interestingly, the viral accessory proteins glycoGag and S2, encoded by murine leukemia virus (MLV) and equine infectious anemia virus (EIAV), respectively, also exhibit SERINC3/5 counteraction activity, despite a lack of homology with Nef (9, 14, 15). Collectively, these studies point to a protective role played by SERINC3/5 during host-retrovirus coevolution over time.

Nef antagonizes the functions of CD4 and MHC class I by sorting them into the endolysosome system. In the case of CD4, Nef uses a di-leucine motif (_164_LL_165_ in the context of ExxxLL) to form a complex with CD4 and AP2, driving the internalization of CD4 into clathrin-coated pits and eventual delivery to the lysosomal compartment (6, 16–18). It has been proposed that the same di-leucine motif of Nef enables AP2-dependent internalization of SERINC3/5, indicating that a single sequence determinant in Nef enables regulation of multiple host factors. It has been suggested that Nef may directly interact with SERINC3/5 to drive its internalization from the cell surface (19–21). However, internalization of SERINC3/5 from the cell surface and exclusion from HIV-1 virions are not prerequisites for Nef-mediated counteraction, indicating that the mechanism by which Nef rescues infectivity remains poorly understood (22). Furthermore, it has been reported that HIV-1 Nef boosts infectivity of viruses produced in certain cell lines in an AP2-dependent even in the absence of SERINC3/5 (23–25). Lastly, Nef variants from Simian Immunodeficiency Viruses (SIV) that fail to antagonize the restriction of HIV-1 by overexpressed human SERINC3/5 retain the ability to enhance HIV-1 infection and spread in human CD4+ T cells (21). These findings point to the existence of unidentified, Nef-sensitive antiviral proteins expressed in human cells that restrict HIV-1 infectivity.

We and others previously demonstrated that interferon-induced transmembrane (IFITM) proteins inhibit HIV-1 infectivity at the stage of fusion, with IFITM3 performing the most potent restriction (26–28). Like SERINC3/5, the anti-HIV activity of IFITM3 is associated with its incorporation into HIV-1 virions and restriction of virion fusion with target cells. However, unlike SERINC3/5, IFITM1-3 are upregulated by interferons (29). We recently showed that, in addition to HIV-1, IFITM3 also restricts the infectivity of MLV. Surprisingly, we found that MLV deficient for glycoGag exhibited heightened sensitivity to restriction by IFITM3, while glycoGag overexpression reduced sensitivity of MLV to IFITM3 (30). Since MLV glycoGag and HIV-1 Nef were previously shown to share the ability to counteract SERINC3/5 and restore retrovirus infectivity, we hypothesized that Nef may also antagonize one or more human IFITM proteins. In further support of an functional relationship between Nef and IFITM proteins, it was recently reported that Nef reduced the levels of IFITM proteins in extracellular vesicles released from T cells, which was caused by downmodulation of IFITM from the cell surface (31). However, the regulation of IFITM proteins by Nef was not explored in the context of HIV-1 infection.

Here we examined whether HIV-1 Nef proteins derived from diverse sources exhibit the ability to counteract the anti-HIV-1 activity of IFITM3 and related human IFITM proteins. We identified primary isolates of Nef that counteracted the loss of HIV-1 infectivity resulting from type-I interferon treatment or IFITM3 overexpression, whereas Nef from lab-adapted molecular clones had little to no effect. Counteraction was AP-2-dependent and correlated to the degree to which the carboxy-terminus of Nef interacted with IFITM3, as measured by co-immunoprecipitation and proximity ligation. Accordingly, IFITM3 counteraction by Nef was associated with reduced cell surface levels of IFITM3 and reduced IFITM3 incorporation into HIV-1 virions. Furthermore, we found that the Nef-IFITM3 interaction impaired IFITM3 dimerization and reduced the impact of IFITM3 on membrane fluidity, which were both previously shown by us to be important for the anti-HIV activity of IFITM3 (32). The antiviral activity of IFITM3 and its negation by Nef were unaffected by SERINC5 knockdown, suggesting that the counteraction of IFITM3 represents a unique function of Nef. By demonstrating that HIV-1 Nef targets an interferon-stimulated gene product, we reveal a previously unrecognized role for HIV-1 Nef in promoting virus infectivity in the setting of the interferon-induced antiviral state.

## RESULTS

To measure the antiviral activity of IFITM3 in HIV-1 producer cells and its sensitivity to Nef, we produced Nef-deficient HIV-1 from HEK293T cells transfected with human IFITM3 (untagged) and individual Nef proteins from diverse sources (tagged with HA). Virus produced was quantified by p24 Gag ELISA and infectivity was measured on TZM-bl reporter cells inoculated with equal quantities of virus. Overexpression of IFITM3 reduced Nef-deficient HIV-1 by approximately 5-fold, but co-expression of IFITM3 with Nef resulted in partial or complete recovery of infectivity depending on the Nef isolate (**Figure 1A and Supplemental Figure 1A**). IFITM3 protein levels following transient transfection of HEK293T were similar to those induced by type-I interferon treatment of the same cell line (**Figure 1B**). Expression of Nef from the lab-adapted molecular clone NL4-3 or a Nef isolate obtained during acute clade H infection from (90CF056) (33) resulted in modestly improved infectivity. In contrast, Nef derived from other acute primary isolates of HIV-1 (93BR020; clade F and 94UG114; clade D) (33) restored HIV-1 infectivity to a greater extent, while a Nef isolate from clade C (97ZA013) fully restored HIV-1 infectivity (**Figure 1A**). We found that 97ZA013 Nef did not significantly boost Nef-deficient HIV-1 infectivity in the absence of IFITM3, while restoration of infectivity in the presence of IFITM3 was a function unique to Nef but not Vpu in the 97ZA013 strain (**Supplemental Figure 1B**). 97ZA013 Nef also counteracted restriction of murine leukemia virus (MLV) by IFITM3 to a similar extent as glycoGag (**Supplemental Figure 1C**). Thus, some HIV-1 Nef proteins exhibit the capacity to counteract the restriction of retrovirus infectivity by IFITM3. Since we and others previously demonstrated that the paralog of IFITM3, IFITM2, also inhibits HIV-1 infectivity (albeit to a lesser extent than IFITM3) (26, 28, 34), we assessed if 97ZA013 Nef regulated the antiviral impacts IFITM2 expression in virus-producing cells. Like IFITM3, restriction of HIV-1 infectivity by IFITM2 was also counteracted by 97ZA013 Nef (**Supplemental Figure 1D**). To determine whether the complete counteraction of IFITM3 by Nef is a conserved feature during early clade C infections, we synthesized a consensus Nef sequence from primary isolates obtained from 105 individuals acutely infected with HIV-1 clade C (35). Similarly to the 97ZA013 isolate, the clade C consensus Nef also fully counteracted the antiviral activity of IFITM3 (**Figure 1A**). Thus, Nef from primary HIV-1 isolates can antagonize human IFITM3, and this is particularly evident for Nef originating from acute clade C, which is the most prevalent form of HIV-1 and accounts for approximately 50% of HIV-1 infections globally (36).

**Figure 1:**
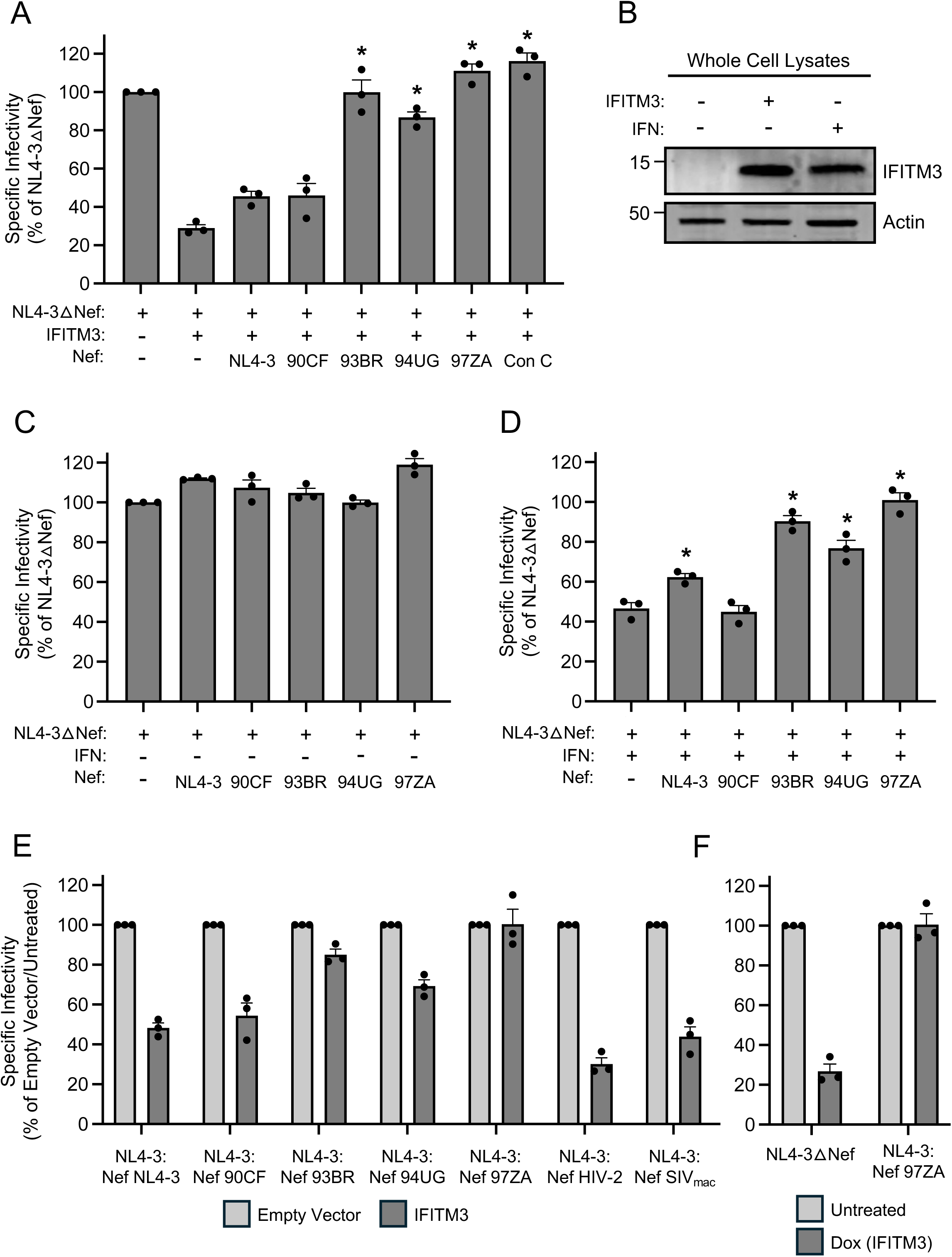
**Nef from diverse HIV-1 group M clades counteract the restriction of HIV-1 infectivity by type-I interferon and IFITM3.** (A) HEK293T cells were co-transfected with NL4-3△Nef (2.0 μg), pCMV-IFITM3 or Empty Vector (0.5 μg), and pBJ-Nef-HA encoding the indicated Nef protein (0.25 μg). Produced virus-containing supernatants were harvested 24 hours post-transfection and quantified by viral p24 ELISA. 25 ng p24 equivalent of fresh virus-containing supernatants were added to TZM-bl cells, and infection was scored by anti-Gag immunostaining at 48 hours post-inoculation. Virus infectivity of each condition is shown as mean and standard deviation (normalized relative to NL4-3△Nef alone, which was set to 100%). Differences that were statistically significant from NL4-3△Nef + IFITM3 as determined by one-way ANOVA are indicated by (*) (*p* < 0.05). (B) HEK293T cells were transfected with pCMV-IFITM3 or Empty Vector (0.5 μg) or treated with type-I interferon (IFN) (∼30 units/well) and whole cell lysates were prepared. SDS-PAGE and immunoblotting were performed with anti-IFITM3 and anti-Actin (used as loading control). Numbers and tick marks left of blots indicate position and size (in kilodaltons) of protein standard in ladder. (C) HEK293T cells were co-transfected with NL4-3△Nef (2.0 μg) and pBJ-Nef-HA encoding the indicated Nef protein (0.25 μg). Produced virus-containing supernatants were harvested 24 hours post-transfection and quantified by viral p24 ELISA. 25 ng p24 equivalent of fresh virus-containing supernatants were added to TZM-bl cells, and infection was scored by anti-Gag immunostaining at 48 hours post-inoculation. Virus infectivity of each condition is shown as mean and standard deviation (normalized relative to NL4-3△Nef alone, which was set to 100%). (D) As in (C), except that transfected cells were treated with type-I IFN overnight prior to harvest of virus-containing supernatants. Virus infectivity of each condition is shown as mean and standard deviation (normalized relative to NL4-3△Nef in (C), which was set to 100%). Differences that were statistically significant from NL4-3△Nef + IFN as determined by one-way ANOVA are indicated by (*) (*p* < 0.05). (E) HEK293T cells were co-transfected with NL4-3 encoding the indicated Nef in cis (2.0 μg) and pCMV-IFITM3 or Empty Vector (0.5 μg). Produced virus-containing supernatants were harvested 24 hours post-transfection and quantified by viral p24 ELISA. 25 ng p24 equivalent of fresh virus-containing supernatants were added to TZM-bl cells, and infection was scored by anti-Gag immunostaining at 48 hours post-inoculation. Virus infectivity of each condition is shown as mean and standard deviation (normalized relative to NL4-3△Nef + Empty Vector, which was set to 100%). (F) SupT1 Tet-ON IFITM3 cells were infected with full-length NL4-3 or NL4-3 encoding 97ZA Nef for 48 hours. Cells were washed and incubated in the presence of doxycycline (500 ng/mL) overnight. Virus-containing supernatants were harvested at 72 hours post-infection (18 hours post-doxycycline) and quantified by viral p24 ELISA. 100 ng p24 equivalent of fresh virus-containing supernatants were added to TZM-bl cells, and infection was scored by anti-Gag immunostaining at 48 hours post-inoculation. Virus infectivity of each condition is shown as mean and standard deviation (normalized relative to Untreated (no doxycycline), which was set to 100%). Filled circles represent biological replicates (independent transfections). Con; consensus. IFN; interferon. Dox; doxycycline.

Since IFITM3 is an interferon-stimulated gene, we next examined whether Nef boosted the infectivity of HIV-1 produced from HEK293T cells treated with type-I interferon. Interferon treatment increased the abundance of IFITM3 protein, but no such increase was observed for its IFITM2 (**Supplemental Figure 1E**). In the absence of interferon treatment, the expression of different Nef constructs had little to no effect on the infectivity of Nef-deficient HIV-1 (**Figure 1C**), suggesting that these Nef proteins do not boost infectivity by counteracting constitutively expressed antiviral factors in HEK293T. However, in cells treated with type-I interferon, certain Nef proteins significantly elevated virus infectivity. Specifically, the three Nef proteins obtained from primary isolates which significantly restored infectivity in cells overexpressing IFITM3 (93BR020, 94UG114, and 97ZA013) also restored infectivity in interferon-treated cells (**Figure 1D**). These data suggest that Nef proteins from certain primary isolates impact the susceptibility of HIV-1 to restriction by interferon-induced antiviral factors including IFITM3.

To confirm our findings on the counteractive role played by certain Nef proteins, we also assessed Nef-mediated counteraction of overexpressed IFITM3 using recombinant, full length HIV-1 encoding different Nef proteins in cis. As was observed transfecting Nef in trans to rescue a Nef-deficient HIV-1, we found that HIV-1 encoding Nef from certain primary isolates (93BR020, 94UG114, and 97ZA013) increased the infectivity of virus produced in the presence of IFITM3, while virus with NL4-3 Nef and 90CF056 Nef did so to a lesser extent (**Figure 1E**). We also found that SIVmac Nef did little to counteract restriction by IFITM3, and HIV-2 Nef was completely inactive in this regard. These data indicate that Nef proteins from primary isolates of HIV-1 can antagonize IFITM3 either when overexpressed or naturally expressed in the context of the provirus.

To measure whether Nef confers resistance to IFITM3 when virus is produced from natural target cells of HIV-1, we measured the infectivity of HIV-1 produced from SupT1 lymphocytes stably expressing IFITM3 under the control of doxycycline (SupT1-IFITM3). In these experiments, SupT1-IFITM3 cells were infected with Nef-deficient HIV-1 or recombinant HIV-1 encoding 97ZA013 Nef and IFITM3 expression was induced or not with doxycycline post-inoculation. Viral supernatants were then collected, and equivalent virus inputs were tested for infectivity in TZM-bl cells. Our results showed that the infectivity of Nef-deficient HIV-1 produced from SupT1-IFITM3 was reduced 5-fold upon IFITM3 induction, while virus encoding 97ZA013 Nef was unaffected by IFITM3 induction (**Figure 1F**). Notably, the amount of ectopic IFITM3 protein induced by doxycycline was similar to the amount of endogenous IFITM3 induced by type-I interferon treatment of SupT1 cells (**Supplemental Figure 1F)**. These findings demonstrate that the antiviral activity of IFITM3 in HIV-producing cells, including T cells, and its resultant impact on virus infectivity, is counteracted by certain Nef proteins derived from primary isolates of HIV-1.

To gain insight into the mechanistic basis for Nef-mediated antagonism of IFITM3, and the basis for antagonism encoded by Nef obtained from primary virus isolates, we compared the effects of two Nef proteins obtained from molecular clones of HIV-1 (SF2 and LAI) and compared them to the activity of 97ZA013 Nef obtained from primary clade C HIV-1 (hereafter referred to as 97ZA). As was observed for NL4-3 Nef (**Figure 1A**), SF2 Nef and LAI Nef modestly boosted HIV-1 infectivity in the presence of IFITM3, while 97ZA Nef fully counteracted IFITM3 (**Figure 2A**). To understand why 97ZA Nef counteracted IFITM3 to a much greater extent than SF2 Nef and LAI Nef, we assessed the degree to which IFITM3 incorporated into HIV-1 virions in their presence. We previously found that restriction of HIV-1 infectivity is associated with the cell surface expression and virion incorporation of IFITM3 oligomers (26, 27, 32). Consistent with its unique capacity to restore infectivity in the presence of IFITM3, the amount of virion-associated IFITM3 protein was reduced by 97ZA Nef (**Figure 2B**). We next tested whether Nef and IFITM3 have the potential to interact following transient transfection in HEK293T cells. In transfected whole cell lysates, we observed that total IFITM3 protein detected by immunoblotting was only modestly affected by the different Nef proteins, with 97ZA Nef slightly decreasing IFITM3 protein (**Figure 2C**). When Nef was immunoprecipitated from lysates, we observed that pull-down of IFITM3 occurred, and 97ZA Nef co-immunoprecipitated with IFITM3 to a greater extent than the other Nefs (**Figure 2D**). Therefore, these data suggest that IFITM3 counteraction by Nef involves a direct or indirect interaction between IFITM3 and Nef, which is associated with partial virion exclusion of IFITM3. We tested the same Nef proteins for counteraction of SERINC5-mediated restriction of HIV-1 infectivity and found that the impact of Nef did not follow the same pattern as was observed with IFITM3. Specifically, SF2 Nef and 97ZA Nef partially counteracted SERINC5 and they did so to a similar magnitude (**Figure 2E**). Thus, the molecular determinants by which Nef counteracts SERINC5 are not identical to those governing counteraction of IFITM3. We excluded that neither the antiviral activity of IFITM3 nor the Nef-mediated boost to infectivity in the presence of IFITM3 were dependent on SERINC5 by performing experiments in SERINC5 knockdown cells (**Supplemental Figure 2A-B**). These findings indicate that antagonism of IFITM3 by Nef from primary isolates is a unique and separate function from that of SERINC5 antagonism. Since we previously demonstrated that IFITM3 reduces HIV-1 entry at the level of membrane fusion with target cells (26), we tested whether the impact of 97ZA Nef on HIV-1 infectivity occurs at the step of virus fusion. Nef-deficient virus was produced and labeled with a self-quenching concentration of DiOC18. Upon addition of equal amounts of labeled virus to TZM-bl cells, DiOC18 de-quenching (fluorescence) in living cells indicated membrane fusion between labeled virus and unlabeled cellular membranes. DiOC18 fluorescence was significantly reduced by IFITM3 in a manner sensitive to 97ZA Nef (**Figure 2F**). These findings suggest that Nef counteracts IFITM3 by interacting with it, partially excluding it from HIV-1 virions, and restoring fusogenic potential to virions.

**Figure 2:**
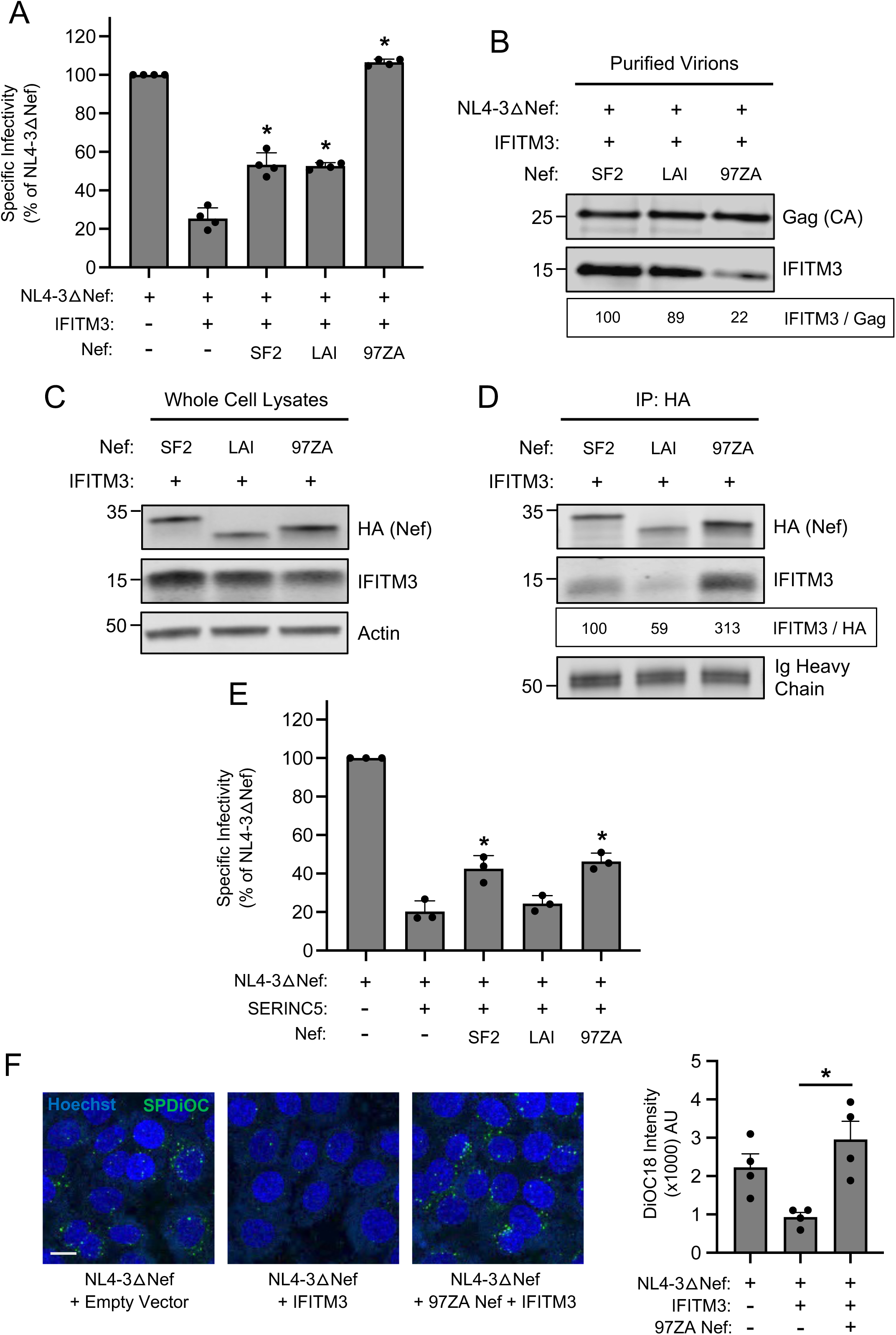
**Nef counteracts IFITM3 and restores HIV-1 entry by interacting with IFITM3 and partially excluding it from virions.** (A) HEK293T cells were co-transfected with NL4-3△Nef (2.0 μg), pCMV-IFITM3 or Empty Vector (0.5 μg), and pBJ-Nef-HA encoding the indicated Nef protein (0.25 μg). Produced virus-containing supernatants were harvested 24 hours post-transfection and quantified by viral p24 ELISA. 25 ng p24 equivalent of fresh virus-containing supernatants were added to TZM-bl cells, and infection was scored by anti-Gag immunostaining at 48 hours post-inoculation. Virus infectivity of each condition is shown as mean and standard deviation (normalized relative to NL4-3△Nef alone, which was set to 100%). Differences that were statistically significant from NL4-3△Nef + IFITM3 as determined by one-way ANOVA are indicated by (*) (*p* < 0.05). (B) Cells were transfected as in (A) and virus-containing supernatants were harvested at 24 hours post-transfection, passed through a 0.45 micron filter, and purified by ultracentrifugation over a 20% sucrose cushion at 25,000 rpm for 1 hour. Viral pellets were subjected to SDS-PAGE and immunoblotting with anti-Gag and anti-IFITM3. Numbers and tick marks left of blots indicate position and size (in kilodaltons) of protein standard in ladder. (C) HEK293T cells were co-transfected with pCMV-IFITM3 or Empty Vector (0.50 μg) and pBJ-Nef-HA encoding the indicated Nef protein (0.25 μg) and whole cell lysates were subjected to SDS-PAGE and immunoblotting with anti-HA, anti-IFITM3, and anti-Actin (which served as loading control). Numbers and tick marks left of blots indicate position and size (in kilodaltons) of protein standard in ladder. (D) From whole cell lysates in (C), Nef proteins were immunoprecipitated with anti-HA followed by SDS-PAGE and immunoblotting with anti-HA and anti-IFITM3. Ig heavy chain served as loading control. Numbers and tick marks left of blots indicate position and size (in kilodaltons) of protein standard in ladder. The IFITM3/HA ratio was calculated for the indicated lanes (normalized to SF2 Nef + IFITM3, which was set to 100%). (E) HEK293T cells were co-transfected with NL4-3△Nef (2.0 μg), pBJ-SERINC5 (0.1 μg) or Empty Vector (0.5 μg), and pBJ-Nef-HA encoding the indicated Nef protein (0.25 μg). Produced virus-containing supernatants were harvested 24 hours post-transfection and quantified by viral p24 ELISA. 25 ng p24 equivalent of fresh virus-containing supernatants were added to TZM-bl cells, and infection was scored by anti-Gag immunostaining at 48 hours post-inoculation. Virus infectivity of each condition is shown as mean and standard deviation (normalized relative to NL4-3△Nef alone, which was set to 100%). Differences that were statistically significant from NL4-3△Nef + SERINC5 as determined by one-way ANOVA are indicated by (*) (*p* < 0.05). Filled circles represent biological replicates (independent transfections). (F) HEK293T cells were co-transfected with NL4-3△Nef (2.0 μg), pCMV-IFITM3 or Empty Vector (0.5 μg), and pBJ-Nef-HA encoding the indicated Nef protein (0.25 μg). Produced virus-containing supernatants were harvested 24 hours post-transfection and one mL was labeled with SP-DiOC18 at a final concentration of 0.2 μM for 60 minutes at room temperature. Labeled virus quantity was measured using p24 ELISA. 40 ng p24 equivalents of labeled virus were added to TZM-bl cells on ice for 1 hour, and then cells were incubated at 37°C for 1 hour and subsequently fixed. Nuclei were stained with Hoechst and fluorescence confocal microscopy was performed. DiOC18 fluorescence intensity in each condition was shown as means and standard deviation. Filled circles represent fields of view containing 8-15 cells each. Scale bar = 10 microns. Differences that were statistically significant between the indicated conditions as determined by one-way ANOVA are indicated by (*) (*p* < 0.05). IP; immunoprecipitation. CA; capsid. AU; arbitrary units.

In order to learn more about the mechanisms employed by Nef to counteract IFITM3, we introduced mutations into 97ZA Nef that have been previously characterized to disrupt various functions of Nef. Compared to 97ZA Nef WT, which fully restored HIV-1 infectivity in the presence of IFITM3, mutation of the myristoylation site of Nef (G2A) resulted in total loss of IFITM3 counteraction (**Figure 3A**). Similarly, mutation of four basic residues (R17/19/21/22A or “R4A4”) in the amino terminus of Nef also prevented counteraction of IFITM3. The G2A and R4A4 mutations are known to disrupt Nef localization to cellular membranes (37–39), indicating that membrane association is an important part of how Nef counteracts IFITM3. To identify the cellular machinery that Nef coopts in order to counteract IFITM3, we introduced L164/165A (“LLAA”) to disrupt the ExxxLL dileucine motif found in the carboxy-terminal loop of Nef, which was previously shown to enable AP-2-dependent downmodulation of CD4 (40–42) and an M20A mutation previously identified to be critical for AP-1-mediated downmodulation of MHC class I (7, 43). The LLAA mutations in Nef resulted in near-complete loss of IFITM3 counteraction, while M20A had no effect (**Figure 3A**). These results indicate that the mechanism used by 97ZA Nef to counteract IFITM3 resembles the mechanism used by Nef to counteract CD4, while it is distinct from the mechanism used to counteract MHC class I. Co-immunoprecipitation experiments revealed that the G2A and R4A4 mutations in Nef inhibited the interaction with IFITM3, while the LLAA and M20A did not (**Figure 3B-C**). Since Nef LLAA exhibits decreased capacity to counteract IFITM3 (**Figure 3A**), yet it still interacts with IFITM3 (**Figure 3C**), the ExxxLL motif likely contributes to IFITM3 counteraction at a step downstream of IFITM3 binding, such as AP-2 recruitment. We also used the proximity ligation assay to confirm that 97ZA Nef WT and IFITM3 interact in intact cells, and with this approach we confirmed that the interaction is disrupted by G2A and R4A4 in Nef (**Figure 3D**). These results suggest that 97ZA Nef counteracts the antiviral activity of IFITM3 and restores virus infectivity by interacting with IFITM3 in membranes and by recruiting AP-2 via the ExxxLL dileucine motif. Since we previously showed that MLV glycoGag counteracted IFITM3, while a glycoGag mutant deficient for AP-2 binding (Y36A) did not (30), these results suggest that glycoGag and Nef use a similar mechanism to counteract IFITM3. To extend those findings and test whether Nef impacts the subcellular localization of IFITM3, we co-transfected HEK293T with IFITM3 and either 97ZA Nef WT or G2A and performed confocal immunofluorescence microscopy. In the absence of Nef, IFITM3 could be detected at the cell surface and intracellular compartments including early endosomes (identified by EEA1-GFP), as demonstrated previously (26, 27, 32) (**Figure 4A**). However, in the presence of 97ZA Nef WT, there was an enrichment of IFITM3 in EEA1-GFP+ puncta, and a portion of Nef protein was found to colocalize with IFITM3 and EEA1-GFP. Furthermore, there was an apparent reduction of IFITM3 from the cell surface. In contrast, the expression of 97ZA Nef G2A, which localized diffusely throughout the cytosol, did not impact the localization of IFITM3 (**Figure 4A**). We quantified the amount of IFITM3 protein on the cell surface by immunostaining living, transfected cells with an anti-IFITM3 antibody, followed by fixation and analysis by flow cytometry. We found that 97ZA Nef WT reduced IFITM3 at the cell surface by roughly 60% (**Figure 4B**). These results are consistent with a mechanistic model whereby Nef uses AP-2 to redirect IFITM3 from the plasma membrane to early endosomes, resulting in decreased virion incorporation of IFITM3 and reduced inhibition of HIV-1 infectivity.

**Figure 3:**
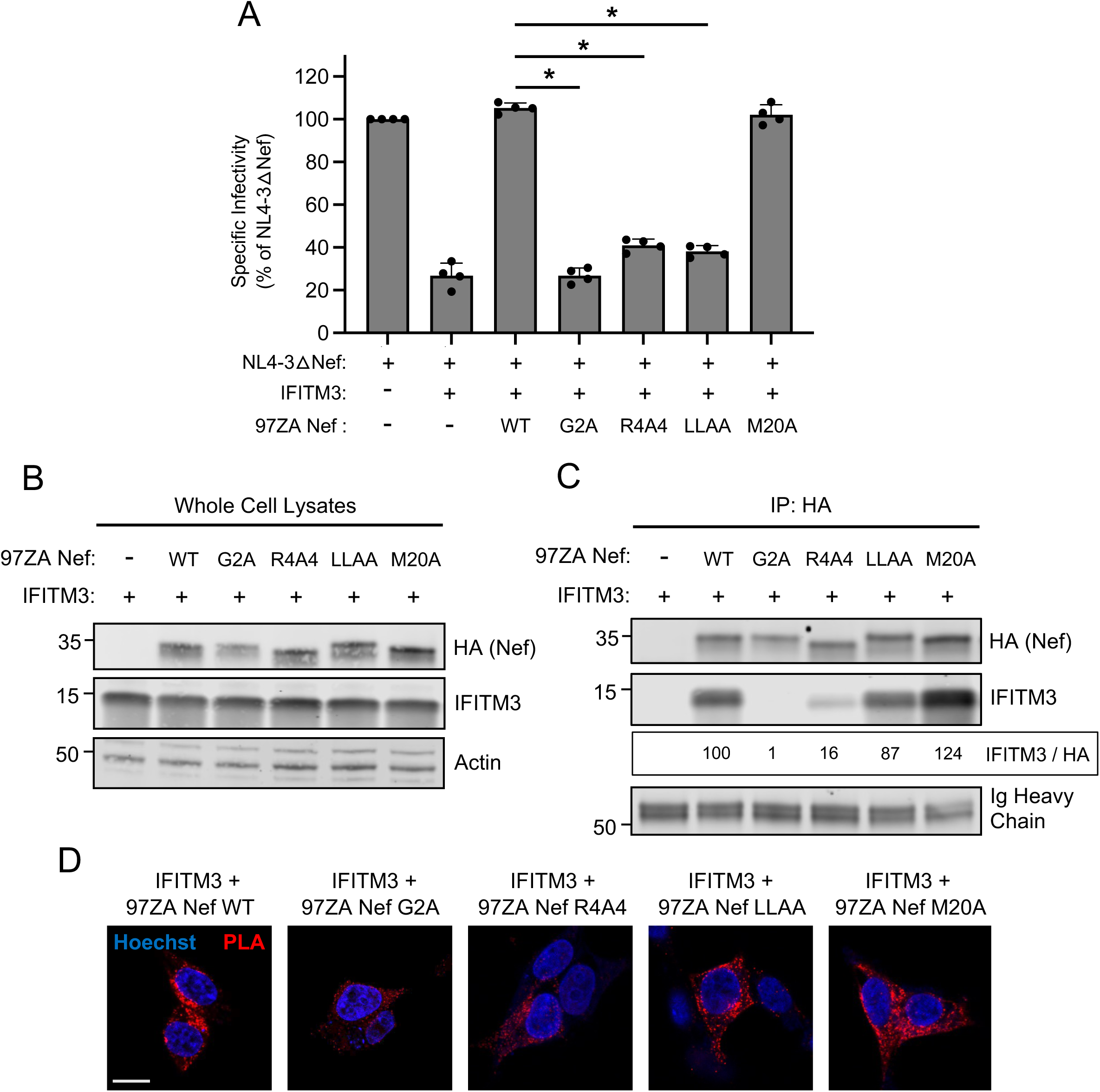
**Nef-mediated counteraction of IFITM3 requires a di-leucine motif known to recruit the endocytic adaptor AP-2.** (A) HEK293T cells were co-transfected with NL4-3△Nef (2.0 μg), pCMV-IFITM3 or Empty Vector (0.5 μg), and pBJ-97ZA Nef-HA encoding WT or mutant protein (0.25 μg). Produced virus-containing supernatants were harvested 24 hours post-transfection and quantified by viral p24 ELISA. 25 ng p24 equivalent of fresh virus-containing supernatants were added to TZM-bl cells, and infection was scored by anti-Gag immunostaining at 48 hours post-inoculation. Virus infectivity of each condition is shown as mean and standard deviation (normalized relative to NL4-3△Nef alone, which was set to 100%). Filled circles represent biological replicates (independent transfections). Differences between the indicated conditions that were statistically significant as determined by one-way ANOVA are indicated by (*) (*p* < 0.05). (B) HEK293T cells were co-transfected with pCMV-IFITM3 or Empty Vector (0.50 μg) and pBJ-97ZANef-HA encoding the WT or mutant (0.25 μg) and whole cell lysates were subjected to SDS-PAGE and immunoblotting with anti-HA, anti-IFITM3, and anti-Actin (which served as loading control). Numbers and tick marks left of blots indicate position and size (in kilodaltons) of protein standard in ladder. (C) From whole cell lysates in (B), Nef proteins were immunoprecipitated with anti-HA followed by SDS-PAGE and immunoblotting with anti-HA and anti-IFITM3. Ig heavy chain served as loading control. Numbers and tick marks left of blots indicate position and size (in kilodaltons) of protein standard in ladder. The IFITM3/HA ratio was calculated for the indicated lanes (normalized to 97ZA Nef WT + IFITM3, which was set to 100%). (D) HEK293T were transfected as in (B), fixed, and proximity ligation assay was performed using anti-IFITM3 and anti-HA followed by confocal microscopy. Nuclei were stained with Hoechst. Scale bar = 10 microns.

**Figure 4:**
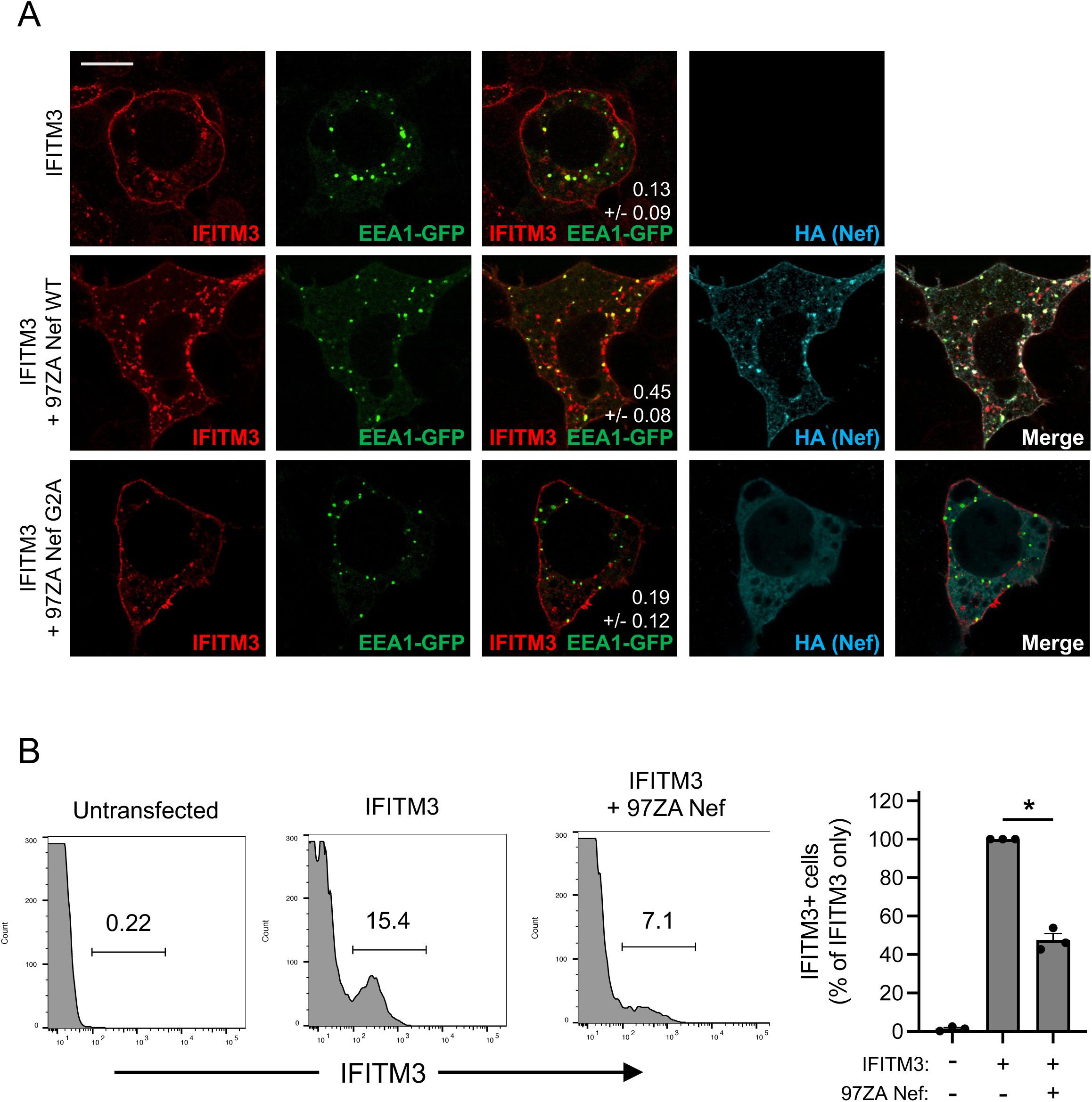
**Nef downmodulates IFITM3 from the cell surface and promotes its accumulation in early endosomes.** (A) HEK293T cells were co-transfected with pCMV-IFITM3 (0.50 μg), pBJ-97ZANef-HA encoding the WT or G2A mutant (0.25 μg), and EEA1-GFP (0.50 μg), fixed, immunostained with anti-IFITM3 and anti-HA, and examined by immunofluorescence confocal microscopy. Pearson’s Correlation Coefficients were calculated between IFITM3 and EEA1-GFP in 20 cells per condition and presented as mean and standard deviation (white numbers in middle column). (B) HEK293T cells were transfected with pCMV-IFITM3 (0.50 μg) alone or both pCMV-IFITM3 and pBJ-97ZANef-HA (0.25 μg) and living, intact cells were stained with anti-IFITM3. Subsequently, cells were fixed, and IFITM3-positive cells were quantified by flow cytometry. A representative example of dot plot histograms is shown on the left and the summary data of three biological replicates (independent transfections) is shown on the right. IFITM3-positive cells in each condition are shown as means and standard deviation (normalized to cells transfected with IFITM3 alone, which was set to 100%). Differences between the indicated conditions that were statistically significant as determined by student’s T test are indicated by (*) (*p* < 0.05). Scale bar = 15 microns.

In addition to partially reducing IFITM3 protein quantity present within virus particles, it remained a possibility that Nef influenced IFITM3 function through additional means. We previously published that oligomerization of IFITM3 is important for its known antiviral functions, including the inhibition of HIV-1 infectivity (32). To measure the degree to which IFITM3 forms oligomers in the absence and presence of Nef, we co-transfected HEK293T with FLAG-tagged IFITM3 as well as Myc-tagged IFITM3. The pull down of Myc-IFITM3 with immunoprecipitated FLAG-IFITM3 demonstrated that these constructs form multimers in transfected cells, as demonstrated previously (32) (**Figure 5A**). However, expression of 97ZA Nef resulted in reduced co-immunoprecipitation of FLAG-IFITM3 and Myc-IFITM3, suggesting that Nef interferes with IFITM3 oligomerization (**Figure 5B**). We and others previously reported that IFITM3 reduces cellular membrane fluidity (32, 44). Furthermore, an oligomerization-defective mutant of IFITM3 exhibited reduced impact on membrane fluidity as well as reduced impact on HIV-1 infectivity (32). Therefore, we measured whether Nef regulated membrane fluidity in cells expressing IFITM3. By measuring membrane fluidity in living cells using fluorescence lifetime imaging of the membrane order probe Flipper-TR (45), we observed that IFITM3 reduced membrane fluidity as expected, as demonstrated by increased Flipper-TR lifetimes (**Figure 5C**). The reduced fluidity observed in IFITM3-expressing cells was reversed using the cholesterol extracting agent methyl-beta-cyclodextrin (MBCD). Notably, 97ZA Nef expression had a similar effect, restoring membrane fluidity to normal levels in IFITM3-expressing cells (**Figure 5C**). These findings suggest that 97ZA Nef not only modifies the subcellular localization of IFITM3, but it also functionally inactivates IFITM3 by inhibiting IFITM3 oligomerization and by inhibiting the effect of IFITM3 oligomers on membrane fluidity. The counteraction through inactivation of IFITM3 by Nef may be a consequence of Nef interacting with IFITM3, either directly or indirectly, in membranes. Therefore, we sought to understand the determinants for IFITM3 binding, and its relationship to IFITM3 counteraction, by producing chimeric Nef proteins. Since NL4-3 Nef and 97ZA Nef differentially counteract IFITM3 (**Figure 1A**), we swapped the carboxy-termini of these Nef proteins and tested them for their ability to rescue Nef-deficient HIV-1 infectivity in the presence of IFITM3. While NL4-3 Nef only modestly counteracted restriction by IFITM3, 97ZA Nef did so fully (**Figure 6A**). However, when the carboxy-terminus of NL4-3 Nef was replaced by that of 97ZA Nef, a complete restoration of HIV-1 infectivity was observed (**Figure 6A**). The reverse swap, whereby the carboxy-terminus of 97ZA Nef was replaced with that of NL4-3 Nef, resulted in a loss of IFITM3 counteraction. These results demonstrate that the carboxy terminus of Nef contains determinants that dictate IFITM3 counteraction. Immunoprecipitation experiments revealed that IFITM3 pulled down to a greater extent with 97ZA Nef relative to NL4-3 Nef, consistent with the greater capacity for 97ZA Nef to counteract IFITM3 functionally (**Figure 6B-C**). Meanwhile, the quantity of IFITM3 pulled down by Nef chimeras indicated that the carboxy-terminus of Nef also governs the interaction with IFITM3 (**Figure 6B-C**). These results suggest that Nef-mediated counteraction of IFITM3 and restoration of HIV-1 infectivity is functionally linked to a physical interaction between IFITM3 and Nef, one that is governed by determinants in the carboxy-terminus of Nef.

**Figure 5:**
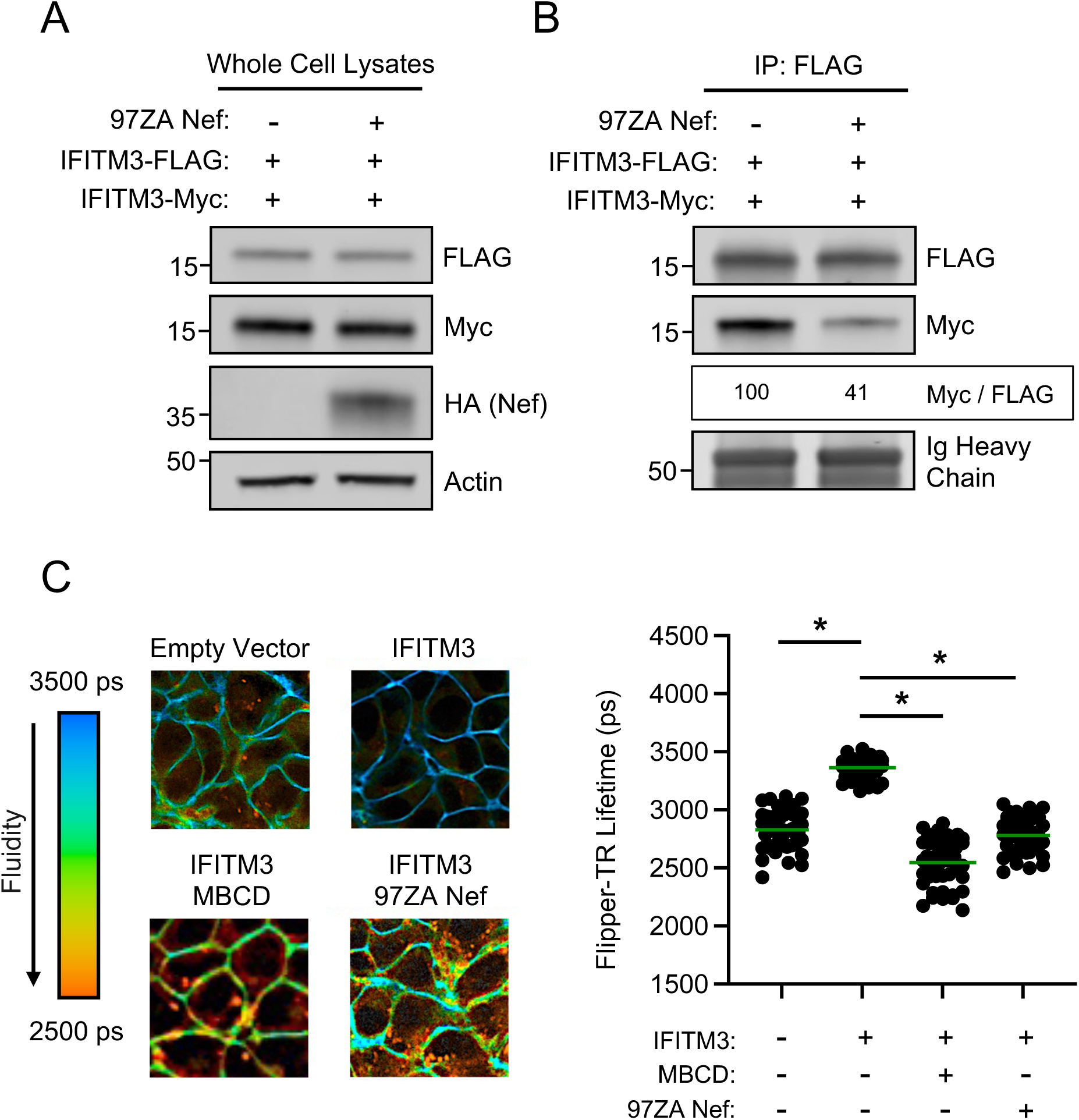
**Nef interferes with IFITM3 oligomerization and restores membrane fluidity in IFITM3-expressing cells.** (A) HEK293T cells were co-transfected with pCMV-IFITM3-FLAG (0.50 μg and pCMV-IFITM3-Myc (0.50 μg) alone or in combination with pBJ-97ZANef-HA (0.25 μg) and whole cell lysates were subjected to SDS-PAGE and immunoblotting with anti-FLAG, anti-Myc, anti-HA, and anti-Actin (which served as loading control). (B) From whole cell lysates in (A), IFITM3-FLAG was immunoprecipitated with anti-FLAG antibody, followed by SDS-PAGE and immunoblotting with anti-FLAG and anti-Myc. Ig Heavy Chain served as loading control. The Myc/FLAG ratio was calculated for the indicated lanes (normalized to IFITM3-FLAG and IFITM3-Myc, which was set to 100%). Numbers and tick marks left of blots indicate position and size (in kilodaltons) of protein standard in ladder. (C) HEK293T cells stably expressing IFITM3 or Empty Vector were untransfected or transfected with pBJ-97ZA Nef-HA (0.25 μg). In the indicated condition, cells were pre-treated with 5 mM MBCD for 2 hours. All conditions were then incubated with the membrane order probe Flipper-TR at a final concentration of 1 μM for 10 minutes. Membrane fluidity of individuals cells was measured by fluorescence lifetime imaging. Left: representative images of fluorescence lifetimes, with blue indicating long lifetimes (less fluid, more rigid) and red indicating short lifetimes (more fluid, less rigid). Right: lifetimes were plotted and mean lifetimes are indicated by green lines, and differences between the indicated conditions that were statistically significant as determined by one-way ANOVA are indicated by (*) (*p* < 0.05). Filled circles correspond to lifetime measurements of individuals cells (50 cells per condition). ps; picoseconds. MBCD; methyl-beta-cyclodextrin.

**Figure 6:**
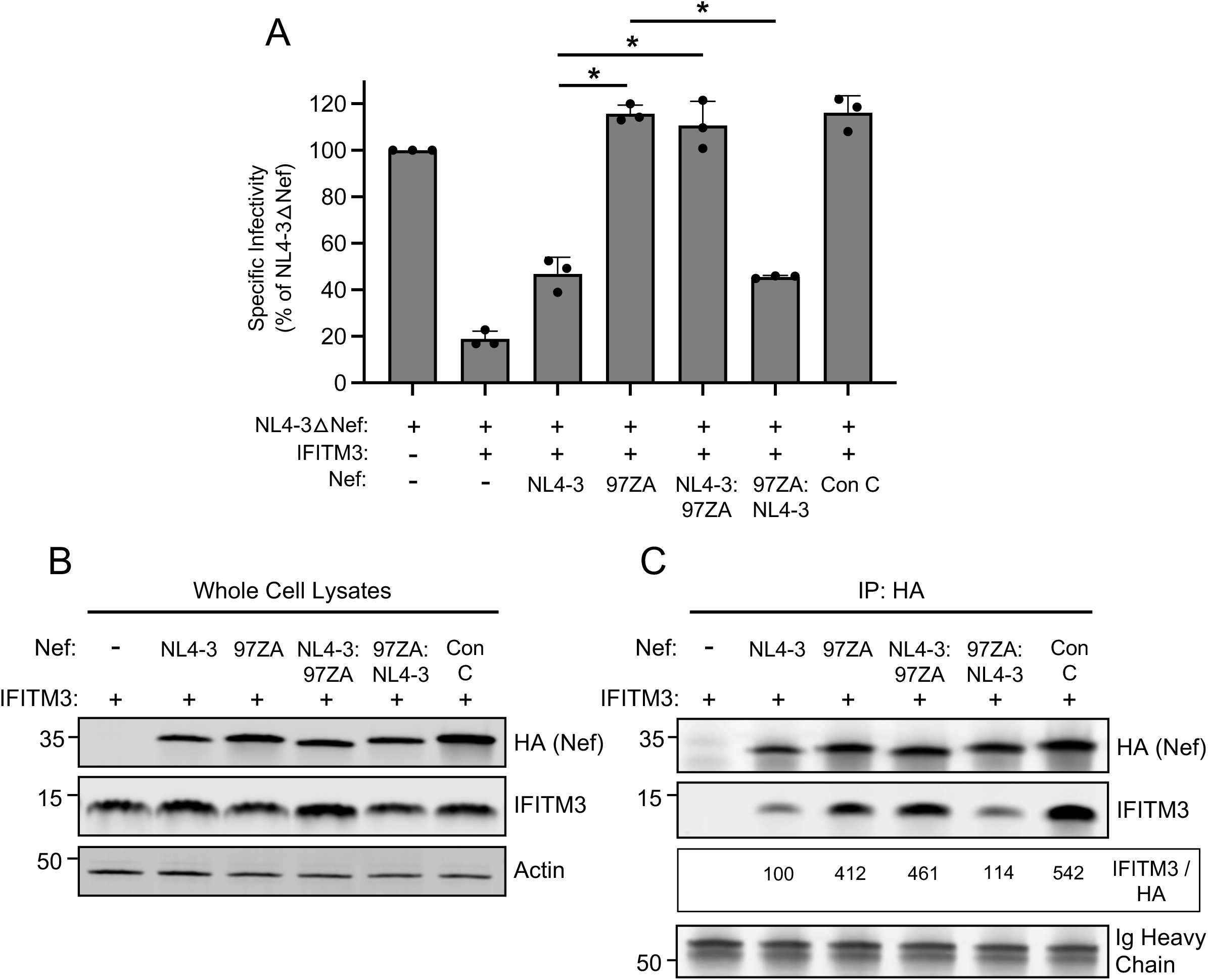
**The carboxy terminus of Nef contains determinants for interacting with and counteracting IFITM3.** (A) HEK293T cells were co-transfected with NL4-3△Nef (2.0 μg), pCMV-IFITM3 or Empty Vector (0.5 μg), and pBJ-NL4-3 Nef-HA, pBJ-97ZA Nef-HA, or chimeric Nef whereby the carboxy termini of NL4-3 and 97ZA Nef proteins were swapped (0.25 μg). Produced virus-containing supernatants were harvested 24 hours post-transfection and quantified by viral p24 ELISA. 25 ng p24 equivalent of fresh virus-containing supernatants were added to TZM-bl cells, and infection was scored by anti-Gag immunostaining at 48 hours post-inoculation. Virus infectivity of each condition is shown as mean and standard deviation (normalized relative to NL4-3△Nef alone, which was set to 100%). Filled circles represent biological replicates (independent transfections). Differences between the indicated conditions that were statistically significant as determined by one-way ANOVA are indicated by (*) (*p* < 0.05). (B) HEK293T cells were co-transfected with pCMV-IFITM3 (0.50 μg) and pBJ-NL4-3 Nef-HA, pBJ-97ZA Nef-HA, or chimeric Nef whereby the carboxy termini of NL4-3 and 97ZA Nef proteins were swapped (0.25 μg), and whole cell lysates were subjected to SDS-PAGE and immunoblotting with anti-HA, anti-IFITM3, and anti-Actin (which served as loading control). (C) From whole cell lysates in (B), Nef proteins were immunoprecipitated with anti-HA and subjected to SDS-PAGE and immunoblotting with anti-HA and anti-IFITM3 (Ig Heavy Chain served as loading control). The IFITM3/HA ratio was calculated for the indicated lanes (normalized to NL4-3 Nef and IFITM3, which was set to 100%). Numbers and tick marks left of blots indicate position and size (in kilodaltons) of protein standard in ladder. Con; consensus. IP; immunoprecipitation.

There exist 20 amino acid differences between the carboxy-termini of NL4-3 Nef and 97ZA Nef (**Figure 7A**). We swapped amino acid residues, individually or in pairs, into 97ZA Nef to assess whether they negatively impacted the counteraction of IFITM3 and its interaction with IFITM3. We found that E153K caused a significant reduction in IFITM3 counteraction by 97ZA Nef, and this was reduced further when combined with S152D (**Figure 7B**). In contrast, the N163T + C164S mutations only somewhat reduced IFITM3 counteraction potential, despite their location in the ExxxLL motif (**Figure 7B**). The others tested (M169V + Q171L and R192F + R193H) did not significantly impact IFITM3 counteraction by 97ZA Nef. Importantly, the mutations that significantly reduced IFITM3 counteraction (S152D + E153K) also significantly impaired IFITM3 pull down (**Figure 7C-D**). Therefore, S152 and E153 in 97ZA Nef are key contributors to the enhanced capacity for this Nef to counteract IFITM3. Furthermore, these results pinpoint key residues in Nef that control binding with IFITM3 and further demonstrate that the interaction between Nef and IFITM3 is pivotal to the counteraction of IFITM3 antiviral activity.

**Figure 7:**
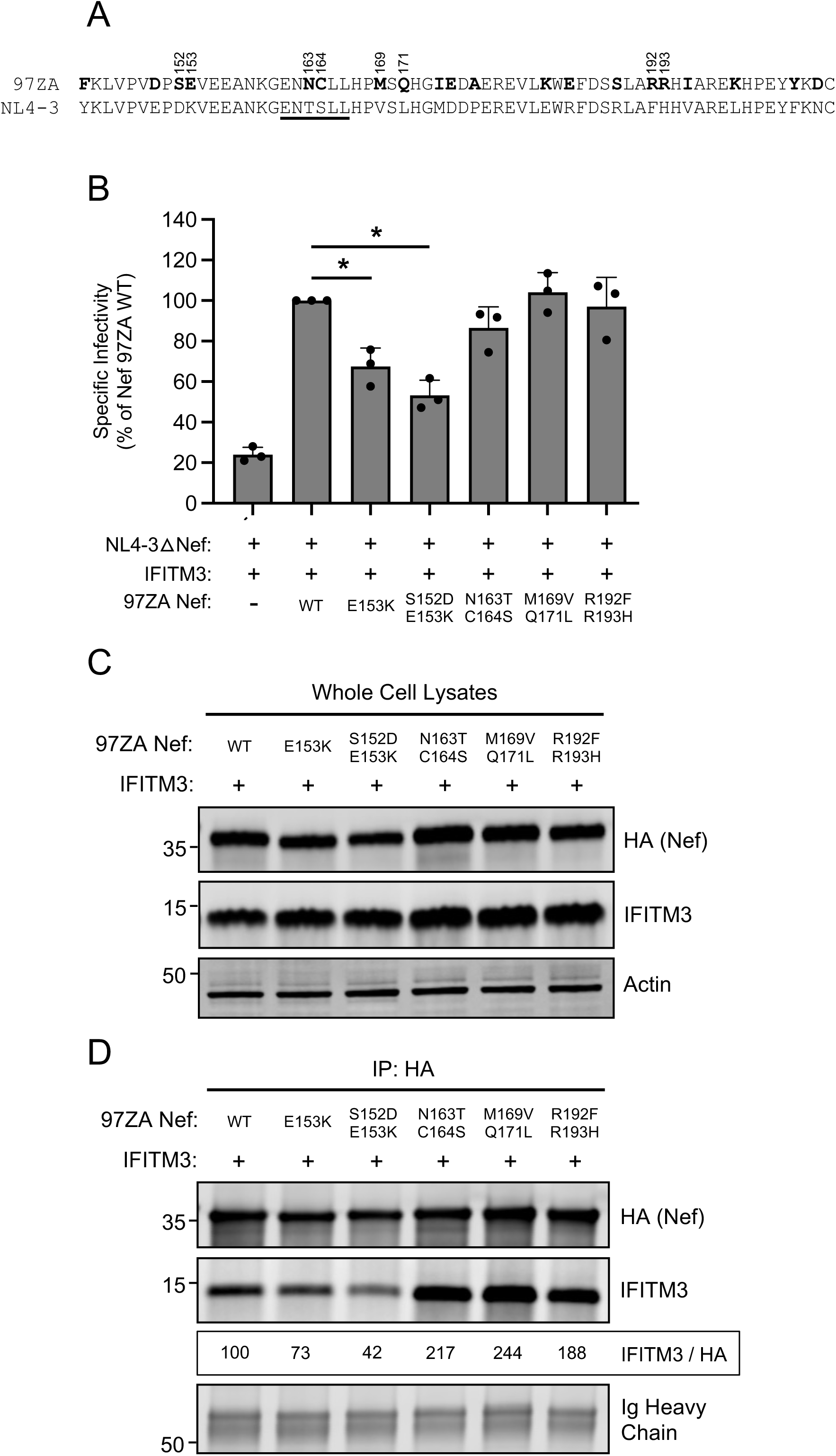
**Residues 152 and 153 in the carboxy terminus of Nef are determinants for interacting with and counteracting IFITM3.** (A) A protein sequence alignment of the carboxy termini (residues 143-206) of 97ZA Nef and NL4-3 Nef. Amino acid residues unique to 97ZA Nef are indicated by bold lettering. The ExxxLL motif previously described to coordinate AP-2 dependent downmodulation of CD4 is underlined. (B) HEK293T cells were co-transfected with NL4-3△Nef (2.0 μg), pCMV-IFITM3 or Empty Vector (0.5 μg), and pBJ-97ZA Nef-HA WT or the indicated mutant (0.25 μg). Produced virus-containing supernatants were harvested 24 hours post-transfection and quantified by viral p24 ELISA. 25 ng p24 equivalent of fresh virus-containing supernatants were added to TZM-bl cells, and infection was scored by anti-Gag immunostaining at 48 hours post-inoculation. Virus infectivity of each condition is shown as mean and standard deviation (normalized relative to NL4-3△Nef + IFITM3 + 97ZA Nef WT, which was set to 100%). Filled circles represent biological replicates (independent transfections). Differences between the indicated conditions that were statistically significant as determined by one-way ANOVA are indicated by (*) (*p* < 0.05). (C) HEK293T cells were co-transfected with pCMV-IFITM3 (0.50 μg) and pBJ-97ZA Nef-HA WT or mutant (0.25 μg), and whole cell lysates were subjected to SDS-PAGE and immunoblotting with anti-HA, anti-IFITM3, and anti-Actin (which served as loading control). (D) From whole cell lysates in (C), Nef proteins were immunoprecipitated with anti-HA and subjected to SDS-PAGE and immunoblotting with anti-HA and anti-IFITM3 (Ig Heavy Chain served as loading control). The IFITM3/HA ratio was calculated for the indicated lanes (normalized to 97ZA Nef and IFITM3, which was set to 100%). Numbers and tick marks left of blots indicate position and size (in kilodaltons) of protein standard in ladder.

Since we found that primary isolates of Nef obtained from acute clade C, clade D, and clade F HIV-1 infection were capable of counteracting HIV-1 restriction by IFITM3 (**Figure 1A**), we next examined Nef proteins from clade B HIV-1, which is the predominant form of HIV-1 in North America. In recent years, much attention has been paid to HIV-1 strains called transmitted/founder (T/F), which represent the phylogenetic ancestors of HIV-1 sequences obtained by single genome amplification from acutely infected individuals (46–48). Interestingly, clade B T/F viruses were previously shown to exhibit elevated infectivity and resistance to interferons relative to viruses obtained during the chronic phase of infection from the same individuals (48). We produced six constructs encoding Nef from the T/F viruses CH040, CH058, CH077, SUMA, TRJO, and WITO and tested them for the ability to rescue IFITM3-restricted HIV-1 infectivity. We found that all six T/F Nef proteins counteracted IFITM3, albeit to different extents. Expression of all Nef constructs were detected in transfected cells, and the heightened activity of CH040 Nef relative to the others was not due to increased Nef protein level (**Supplemental Figure 3**). Nef proteins from CH040, CH058, and CH077 rescued HIV-1 infectivity to the greatest extent, with CH040 completely restoring HIV-1 infectivity (**Figure 8A**). In comparison, the other T/F Nef proteins partially restored infectivity. We also tested full length CH040 virus for sensitivity to restriction by IFITM3, and we found that it was resistant compared to full length NL4-3 (**Figure 8B**). Using additional constructs in which Nef proteins were appended with an HA tag (for the purpose of enabling immunoprecipitation), we confirmed that CH040 Nef completely counteracts IFITM3 while SUMA Nef does so to an intermediate extent (**Figure 8C**). To determine whether these two T/F Nef proteins differ with respect to the IFITM3 interaction, we performed immunoprecipitation of Nef and assess IFITM3 protein pull down. In accordance with its improved counteraction of IFITM3, CH040 Nef pulled down more IFITM protein than SUMA Nef (**Figure 8D-E**). Overall, in addition to the single representatives of HIV-1 clade D, clade F, clade C, and the clade C consensus, at least some clade B T/F viruses encode Nef proteins capable of interacting with and antagonizing the antiviral activity of IFITM3.

**Figure 8:**
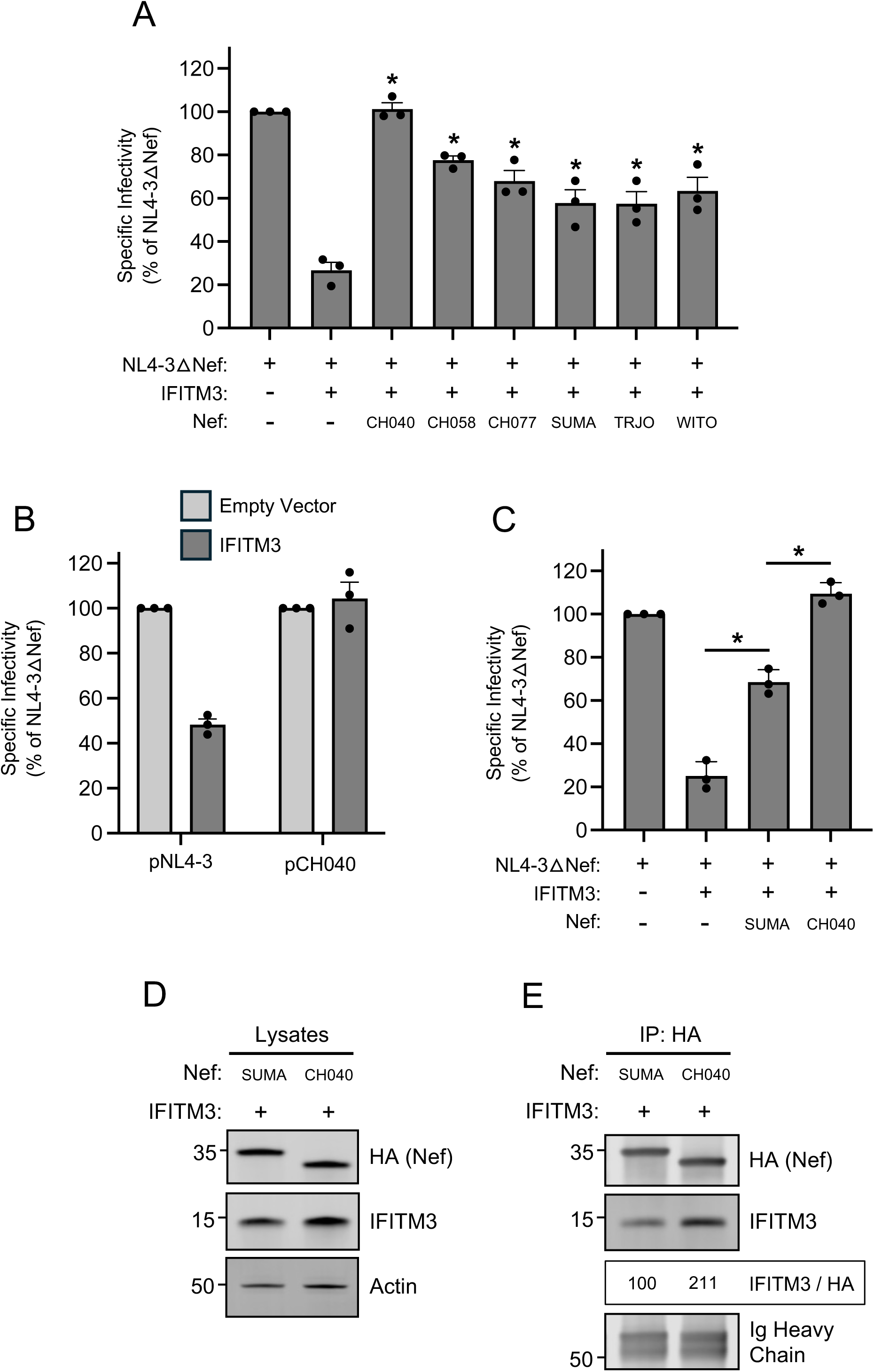
**Nef from HIV-1 clade B T/F viruses counteract restriction by IFITM3.** (A) HEK293T cells were co-transfected with NL4-3△Nef (2.0 μg), pCMV-IFITM3 or Empty Vector (0.5 μg), and pCl-Nef encoding the indicated Nef protein (0.25 μg). Produced virus-containing supernatants were harvested 24 hours post-transfection and quantified by viral p24 ELISA. 25 ng p24 equivalent of fresh virus-containing supernatants were added to TZM-bl cells, and infection was scored by anti-Gag immunostaining at 48 hours post-inoculation. Virus infectivity of each condition is shown as mean and standard deviation (normalized relative to NL4-3△Nef alone, which was set to 100%). Filled circles represent biological replicates (independent transfections). Differences between NL4-3△Nef + IFITM3 and the indicated conditions that were statistically significant as determined by one-way ANOVA are indicated by (*) (*p* < 0.05). (B) HEK293T cells were co-transfected with full length pNL4-3 or full length pCH040 (2.0 μg) and pCMV-IFITM3 or Empty Vector (0.5 μg). Produced virus-containing supernatants were harvested 24 hours post-transfection and quantified by viral p24 ELISA. 25 ng p24 equivalent of fresh virus-containing supernatants were added to TZM-bl cells, and infection was scored by anti-Gag immunostaining at 48 hours post-inoculation. Virus infectivity of each condition is shown as mean and standard deviation (normalized relative to Empty Vector, which was set to 100%). (C) HEK293T cells were co-transfected with NL4-3△Nef (2.0 μg), pCMV-IFITM3 or Empty Vector (0.5 μg), and pBJ-Nef-HA encoding the indicated Nef protein (0.25 μg). Produced virus-containing supernatants were harvested 24 hours post-transfection and quantified by viral p24 ELISA. 25 ng p24 equivalent of fresh virus-containing supernatants were added to TZM-bl cells, and infection was scored by anti-Gag immunostaining at 48 hours post-inoculation. Virus infectivity of each condition is shown as mean and standard deviation (normalized relative to NL4-3△Nef alone, which was set to 100%). Filled circles represent biological replicates (independent transfections). Differences between NL4-3△Nef + IFITM3 and the indicated conditions that were statistically significant as determined by one-way ANOVA are indicated by (*) (*p* < 0.05). (D) HEK293T cells were co-transfected with pCMV-IFITM3 (0.50 μg) and pBJ-Nef-HA encoding the indicated Nef protein (0.25 μg), and whole cell lysates were subjected to SDS-PAGE and immunoblotting with anti-HA, anti-IFITM3, and anti-Actin (which served as loading control). (E) From whole cell lysates in (D), Nef proteins were immunoprecipitated with anti-HA and subjected to SDS-PAGE and immunoblotting with anti-HA and anti-IFITM3 (Ig Heavy Chain served as loading control). The IFITM3/HA ratio was calculated for the indicated lanes (normalized to SUMA Nef and IFITM3, which was set to 100%). Numbers and tick marks left of blots indicate position and size (in kilodaltons) of protein standard in ladder.

## DISCUSSION

Here we report that restriction of HIV-1 infectivity following ectopic IFITM3 expression in virus-producing cells (epithelial cells or T lymphocytes) is counteracted by Nef proteins obtained from primary isolates of acute HIV-1 infection. In particular, we find that Nef proteins obtained from individuals acutely infected with HIV-1 clade C, the predominant form of circulating HIV-1 worldwide, are especially potent counteractors of IFITM3. Furthermore, Nef proteins that counteracted ectopic IFITM3 also counteracted the antiviral state elicited by type-I interferon treatment. Using clade C Nef to uncover the mechanistic basis for IFITM3 counteraction, we found that Nef interacts with IFITM3, utilizes an AP-2-dependent pathway to downmodulate IFITM3 from the cell surface, and partially excludes IFITM3 from HIV-1 virions. Furthermore, Nef inhibits IFITM3 oligomerization and reduces the impact of IFITM3 oligomers on cellular membrane fluidity, indicating that, in addition to altering the subcellular localization of IFITM3, Nef also functionally inactivates IFITM3. Our findings may suggest that counteraction of IFITM3 by Nef is important during early stages of HIV-1 infection in vivo. This notion is supported by our findings that Nef proteins from clade B T/F viruses also exhibit the capacity to antagonize IFITM3. It will be important to assess whether counteraction of IFITM3 is also a feature of Nef proteins isolated during chronic stages of HIV-1 infection. Nonetheless, it is possible that this previously uncharacterized activity of Nef is particularly important in the context of the interferon-induced antiviral state, for example, during the initial infection of and virus spread within mucosal tissue following virus transmission.

While performing the experiments detailed in this manuscript, it was reported that HIV-1 Nef reduces the levels of IFITM proteins found in extracellular vesicles released from T cells (31). Mechanistically, Nef downmodulated constitutively expressed human IFITM1, IFITM2, and IFITM3 from the surface of T cells and reduced IFITM1-3 levels in detergent-resistant lipid rafts. Importantly, those results were obtained using a primary isolate of Nef (31). Our findings presented here, in the context of HIV-1 infection, corroborate those data in that primate isolates of Nef counteracted IFITM3 antiviral function in a manner associated with its downmodulation from the cell surface. While the majority of our work was performed using HEK293T epithelial cells as virus-producing cells, we also found that Nef countered ectopic IFITM3 and boosted HIV-1 infectivity when employing a Dox-inducible SupT1 lymphocyte cell line. It will be of interest to address whether Nef influences the localization of IFITM3 to lipid rafts in virus-producing cells. Indeed, early studies on human IFITM proteins implicated them as part of signaling complexes associated with lipid rafts (49, 50). Palmitoylation is a post-translational modification that promotes the localization of transmembrane proteins to lipid rafts (51). Human IFITM3 is palmitoylated at conserved cysteine residues, and this lipidation promotes membrane anchoring and promotes antiviral functions (27, 52, 53) including the restriction of HIV-1 (27). IFITM3 also directly interacts with membrane cholesterol (54–56), and its inhibition of membrane fluidity may depend upon cholesterol (57). Interestingly, Nef was reported to be a cholesterol-binding, lipid raft-localizing protein and these traits are associated with its ability to promote HIV-1 infectivity (58). Since egress from cholesterol-enriched lipid rafts in virus-producing cells is important for HIV-1 infectivity (59–63), is possible that raft-associated IFITM3 reduces the level of cholesterol available to budding virions while Nef restores cholesterol levels there, possibly by excluding IFITM3 from rafts.

HIV-1 was previously reported to boost HIV-1 infectivity by counteracting the antiviral activities of SERINC3/5, which are not regulated by interferons (9, 10). In that sense, our demonstration that primary isolates of Nef can counteract IFITM3 may represent a rare instance of HIV-1 Nef targeting an interferon-stimulated gene product. HIV-1 Nef has been reported to regulate the production of interferon by interfering with STAT1 phosphorylation (14), but less is known about the direct antagonism of interferon-stimulated effector proteins by Nef. Another HIV-1 accessory protein, Vpu, antagonizes the interferon-inducible protein BST2/tetherin (64, 65), but in certain non-human lentiviruses that lack a Vpu open reading frame, such as Simian Immunodeficiency Viruses, antagonism of tetherin is performed by Nef (66, 67). Furthermore, it was shown that certain isolates of HIV-1 group M encode Nef proteins with the functional capacity to counteract human tetherin (68). Therefore, our work here indicates that IFITM3 can be added to the short list of interferon-stimulated gene products targeted by HIV-1 Nef.

Our results suggest that Nef may boost HIV-1 infectivity by targeting multiple cellular factors, including IFITM3, that restrict HIV-1 infectivity. However, it is worth noting that the mechanisms used by Nef co counteracts SERINC3/5 and IFITM3 are similar yet distinct. This is demonstrated by the sensitivity of SERINC3/5 to different Nef proteins as compared to the sensitivity of IFITM3 to those same Nef proteins. It was previously shown that Nef from the clade H isolate 90CF056 poorly counteracts SERINC3/5 (10, 69), while Nef from clade C isolate 97ZA is a more potent antagonist (10). This matches the effects of those same Nef proteins on IFITM3-mediated restriction (**Figure 1A**). However, 97ZA Nef completely restores HIV-1 infectivity in the presence of IFITM3 but only partially restores infectivity in the presence of SERINC5 (**Figure 2E**). Furthermore, SF2 Nef counteracts SERINC5 to a similar extent as 97ZA Nef (**Figure 2E**)(10), whereas these Nef proteins differentially counteract IFITM3 (**Figure 2A**). These results indicate that the sequence determinants used by Nef to counteract SERINC3/5 versus those used to counteract IFITM3 are non-identical. We found that the carboxy terminus of Nef dictates the degree to which both IFITM3 interaction and IFITM3 counteraction occur (**Figure 6A-C**). More specifically, we identified the carboxy-terminal residues S152 and E153 as being highly consequential for IFITM3 binding and its counteraction (**Figure 7A-C**). Future work will highlight the residues in IFITM3 that are targeted by the carboxy terminus of Nef in order to gain a greater understanding of the IFITM3-Nef binding interface, which may involve other cellular proteins like AP-2. In that regard, recombinant IFITM3, Nef, and AP-2 proteins could be used to measure the direct binding potential between these proteins, and structural characterization may help illuminate the protein-protein and protein-protein-protein interfaces involved. Furthermore, it will be interesting and important to establish the species-specific nature (or lack thereof) of the IFITM3-Nef relationship. For example, determining whether non-human IFITM3 proteins are sensitive to HIV-1 Nef and whether human IFITM3 is sensitive to Nef from other primate lentiviruses may reveal additional details about the IFITM3-Nef functional interface, and will determine whether coevolution occurs at this host-virus interaction as part of an ongoing genetic conflict.

---

**Supplemental Figure 1:**

(A) HEK293T cells were co-transfected with pCMV-IFITM3 (0.50 μg) and pBJ-Nef-HA encoding the indicate Nef protein (0.25 μg), and whole cell lysates were subjected to SDS-PAGE and immunoblotting with anti-HA, anti-IFITM3, and anti-Actin (which served as loading control). Numbers and tick marks left of blots indicate position and size (in kilodaltons) of protein standard in ladder. (B) HEK293T cells were co-transfected with NL4-3△Nef (2.0 μg), pCMV-IFITM3 or Empty Vector (0.5 μg), and pBJ-97ZA Nef-HA or pBJ-97ZA Vpu-HA (0.25 μg). Produced virus-containing supernatants were harvested 24 hours post-transfection and quantified by viral p24 ELISA. 25 ng p24 equivalent of fresh virus-containing supernatants were added to TZM-bl cells, and infection was scored by anti-Gag immunostaining at 48 hours post-inoculation. Virus infectivity of each condition is shown as mean and standard deviation (normalized relative to NL4-3△Nef alone, which was set to 100%). Filled circles represent biological replicates (independent transfections). (C) HEK293T cells were co-transfected with MLV△glycoGag (2.5 μg), pBabeLuc (0.6 μg), pCMV-Xenogp85 (xenotropic Env) (0.5 μg), and pCMV-IFITM3 or Empty Vector (0.5 μg), and where indicated, pCMV-glycoGag-Myc (0.25 μg) or pBJ-97ZA Nef-HA (0.25 μg). Produced virus-containing supernatants were harvested 48 hours post-transfection and quantified by viral Gag immunoblotting with anti-p30 of pelleted viruses. Equal volumes of p24 equivalent of fresh virus-containing supernatants were added to HT1080-mCAT1cells, and infection was scored by luciferase assay at 48 hours post-inoculation. Luciferase values were divided by Gag immunoblot intensity to derive a specific infectivity measurement. Virus infectivity of each condition is shown as mean and standard deviation (normalized relative to MLV△glycoGag alone, which was set to 100%). Differences that were statistically different from the indicated condition and MLV△glycoGag + IFITM3 are indicated by (*) (*p* < 0.05). Filled circles represent biological replicates (independent transfections). (D) HEK293T cells were co-transfected with NL4-3△Nef (2.0 μg), pCMV-IFITM3 or pCMV-IFITM2 or Empty Vector (0.5 μg), and pBJ-97ZA Nef-HA (0.25 μg). Produced virus-containing supernatants were harvested 24 hours post-transfection and quantified by viral p24 ELISA. 25 ng p24 equivalent of fresh virus-containing supernatants were added to TZM-bl cells, and infection was scored by anti-Gag immunostaining at 48 hours post-inoculation. Virus infectivity of each condition is shown as mean and standard deviation (normalized relative to NL4-3△Nef alone, which was set to 100%). Filled circles represent biological replicates (independent transfections). (E) HEK293T cells were untreated or treated with ∼30 units type-I IFN (IFN Beta 1a) for 18 hours and whole cell lysates were subjected to SDS-PAGE and immunoblotting with anti-IFITM3, anti-IFITM2, and anti-Actin (which served as loading control). Numbers and tick marks left of blots indicate position and size (in kilodaltons) of protein standard in ladder. (F) SupT1 Tet-On IFITM3 (SupT1-IFITM3) cells were untreated, treated with doxycycline (500 ng/mL), or treated with type-I IFN (IFN Beta 1a) (∼30 units) for 18 hours. Cells were fixed and immunostained with anti-IFITM3 antibody and analyzed with flow cytometry. Unstained cells served as background fluorescence.

**Supplemental Figure 2:**

(A) Stable SERINC5 knockdown was performed in HEK293T cells as previously described, and knockdown was assessed by quantitative RT-PCR (70). SERINC5 mRNA levels per 1x10^5 copies of GAPDH mRNA were plotted. (B) SERINC5 knockdown HEK293T cells were co-transfected with NL4-3△Nef (2.0 μg), pCMV-IFITM3 or Empty Vector (0.5 μg), and pBJ-97ZA Nef-HA (0.25 μg). Produced virus-containing supernatants were harvested 24 hours post-transfection and quantified by viral p24 ELISA. 25 ng p24 equivalent of fresh virus-containing supernatants were added to TZM-bl cells, and infection was scored by anti-Gag immunostaining at 48 hours post-inoculation. Virus infectivity of each condition is shown as mean and standard deviation (normalized relative to NL4-3△Nef alone, which was set to 100%). Differences between the indicated conditions that were statistically significant as measured by student’s T test are indicated by (*) (*p* < 0.05). Filled circles represent biological replicates (independent transfections).

**Supplemental Figure 3:**

HEK293T cells were co-transfected with pCMV-IFITM3 (0.50 μg) and pCl-Nef encoding the indicate Nef protein (0.25 μg), and whole cell lysates were subjected to SDS-PAGE and immunoblotting with anti-Nef, anti-IFITM3, and anti-Actin (which served as loading control). Numbers and tick marks left of blots indicate position and size (in kilodaltons) of protein standard in ladder.

## METHODS

### Cells and cell culture reagents

HEK293T (ATCC; CRL-3216), TZM-bl (NIH HIV Reagents Program/BEI; HRP-8129) and HT1080-mCAT1 (described previously (71)) were cultured in Dulbecco’s modified Eagle’s medium (DMEM, Gibco) supplemented with 10% heat-inactivated fetal bovine serum (FBS, Hyclone) and 1% penicillin-streptomycin (Gibco) at 37°C with 5% CO_2_. HEK293T stably expressing Empty Vector and IFITM3 were previously described (32). Tet ON SupT1-IFITM3 cells were previously described (26, 72) and were cultured in RPMI (Gibco) supplemented with 10% heat-inactivated fetal bovine serum (FBS, Hyclone) and 1% penicillin-streptomycin (Gibco) at 37°C with 5% CO_2_. Doxycycline hyclate was obtained from Sigma (D5207). Human interferon beta 1a was obtained from PBL Assay Science (11415–1).

### Plasmids and molecular biology

pCMV-hIFITM3, pCMV-hIFITM2, and Empty Vector were obtained from Origene. pBJ-97ZA Nef-HA, pBJ-90CF Nef-HA, pBJ-93BR Nef-HA, pBJ-94UG Nef-HA, pBJ-SF2 Nef-HA, pBJ-LAI Nef-HA, and pBJ-SERINC5 were obtained from Heinrich Gottlinger (University of Massachusetts). pCl-CH040 Nef, pCl-THRO Nef, pCl-WITO Nef, pCl-CH077 Nef, pCl-CH058 Nef, pCl-TRJO Nef, and pCl-SUMA Nef were obtained from Mary Lewinski and John Guatelli (University of California San Diego). pBJ-NL4-3 Nef-HA, pBJ-CH040 Nef-HA, pBJ-SUMA Nef-HA, chimeric pBJ-NL4-3/97ZA Nef-HA and pBJ-97ZA/NL4-3 Nef-HA, and mutants of pBJ-97ZA Nef-HA were produced in this study by PCR amplification and BamH1/EcoR1 restriction sites. A consensus Nef derived from 105 acute HIV-1 clade C infections (35) was generated online (https://www.hiv.lanl.gov/content/sequence/CONSENSUS/SimpCon.html) and cloned into pBJ to produce the pBJ-Con C Nef-HA construct. The following full length virus constructs, and their sources, are as follows: pNL4-3 (Olivier Schwartz, Pasteur Institute), NL4-3△Nef (Olivier Schwartz, Pasteur Institute), pCH040 (Nicoletta Casartelli and Olivier Schwartz, Pasteur Institute), pNL4-3 encoding 90CF, 93BR, 94UG, and 97ZA Nef (Heinrich Gottlinger, University of Massachusetts), pNL4-3 encoding HIV-2 BEN Nef or SIVmac239 Nef (73)(Richard Sloan, University of Edinburgh). The plasmids encoding glycoGag-deficient Moloney MLV Gag-Pol, pCMV-glycoGag-Myc, pBabeLuc, and pCMV-Xenogp85 (xenotropic Env) were previously described (30). EEA1-GFP was obtained from Addgene (42307). A list of Nef protein sequences used in this study is found in **Supplemental File 1**.

### Virus production and infections

For virus production in HEK293T, cells were seeded (200,000 cells/well) in a 6-well plate and transfected with the indicated viral plasmids using Mirus TransIT-293 transfection reagent (Mirus Bio; 2700). Virus containing supernatants were harvested at 24 hours post-transfection and centrifuged at 2500 rpm for 5 min to remove cellular debris. HIV-1 quantity was measured using the HIV-1 p24 ELISA Kit (XpressBio; XB-1000) while MLV quantity was measured using anti-p30 immunoblotting. To measure specific HIV-1 infectivity, 25 ng p24 equivalents were added to TZM-bl cells. Cells were fixed/permeabilized with Cytofix/Cytoperm Solution (BD; 554722) at 48 hours post-infection and immunostained with anti-Gag KC57-FITC (Beckman Coulter; 6604665). Samples were acquired on a BD LSRFortessa flow cytometer and analyzed using FlowJo software (version 10.8.1). To measure MLV infectivity, virus was added to HT1080-mCAT1 cells and infection was measured at 48 hours post-infection using luciferase assay (Promega; E1501).

For virus production in Tet-ON SupT1-IFITM3 cells, 10,000 cells were seeded in a 48-well plate and inoculated with 200 ng p24 equivalents of NL4-3△Nef or NL4-3(97ZA Nef). 18 hours later, cells were washed with PBS and resuspended in fresh RPMI containing Doxycycline (500 ng/mL) or not. At 3 days Dox addition, cells were spun down, virus-containing supernatants were harvested, and virus was quantified by p24 ELISA. 75 ng p24 equivalents were added to TZM-bl cells, and cells were fixed at 48 hours post-infection and immunostained with KC57-FITC to enumerate infection by flow cytometry.

For virus entry/fusion measurements, HIV-1 produced from HEK293T cells was labeled with SP-DiOC18 (Thermo Fisher; D7778) at a final concentration of 0.2 μM. Labeled viruses were filtered through 0.22 micron filters to remove virus aggregates and virus quantity was measured using HIV-1 p24 ELISA Kit (XpressBio, XB-1000). 40 ng p24 equivalents of labeled virus was added to TZM-bl cells seeded (80,000 cells/well) in an 8-well Mu slide dish (Ibidi; 80826), and cells were incubated for 1 hour on ice. Cells were washed three times with serum-free DMEM, fresh, serum-containing DMEM was added to cells, and cells were incubated for 1 hour at 37°C.

Cells were then fixed with 4% formaldehyde for 15 mins at 4°C. Nuclei were labeled with Hoechst 33342 (Thermo Fisher; 62249) for 10 mins at room temperature. SP-DiOC18 fluorescence was measured imaged on a Leica Stellaris confocal fluorescence microscope. Images were analyzed and SP-DiOC18 fluorescence intensities were measured in Fiji (ImageJ).

### Flow cytometry

For quantification of HIV-1 infection, TZM-bl cells were fixed with Cytofix/Cytoperm Solution (BD; 554722) and stained with Anti-HIV-1 Core Antigen KC57-FITC (Beckman Coulter; 6604665) for 30 mins at room temperature.

For cell surface staining of IFITM3, transfected HEK293T were stained with anti-IFITM3 (Proteintech; 66081-1) on ice for 30 mins and then fixed with 2% formaldehyde solution (Invitrogen; FB002) for 10 mins at room temperature. Cells were then washed with PBS and stained with secondary antibody goat anti-mouse IgG (H+L) Alexa Fluor 488 (Invitrogen; A11001).

For staining of total IFITM3 in Tet ON SupT1-IFITM3 cells, cells were fixed with Cytofix/Cytoperm Solution (BD; 554722) and stained with anti-IFITM3 (Abcam; ab109429). Cells were washed with Cyto Perm/Wash buffer (BD; 554723) and stained with goat anti-rabbit IgG (H+L) Alexa Fluor 647.

Cells were analyzed on a BD LSRFortessa flow cytometer and analyzed using FlowJo software (versions 10.8.1).

### Western blot analysis

Whole cell lysis was performed using a buffer consisting of 20 mM HEPES, 150 mM NaCl, 1 mM EDTA, and 1% Triton X-100 (Sigma; X100) containing Halt protease inhibitor cocktail, EDTA-free (Thermo Fisher; 78425). Lysis was performed on ice for 30 mins prior to centrifugation at 12,000 rpm for 10 mins at 4°C, and supernatants were harvested. Protein quantity was measured using the Protein Assay (Bio-Rad; 5000001). Lysates were mixed with NuPAGE Reducing Agent (Invitrogen; NP0009) and Protein Loading Buffer (Li-COR; 928-40004) and loaded into Criterion XT 12% polyacrylamide Bis-Tris gels (Bio-Rad; 3450117). SDS-PAGE was performed with NuPAGE MES SDS Running Buffer (Invitrogen; NP0002). Proteins were transferred to Immobilion PVDF membrane, pore size 0.45 micron (Millipore; IPFL00010). Membranes were blocked with Intercept PBS Blocking Buffer (Li-COR; 927-70001) for 30 mins at room temperature. The following primary antibodies were used: anti-IFITM3 (Abcam; ab109429), anti-IFITM2 (Proteintech; 66137-1-Ig), anti-HA (Abcam; ab9110), anti-Actin (Santa Cruz Biotechnology; sc-47778), anti-Nef (NIH HIV Reagent Program/BEI; ARP-2949), anti-Gag (183-H12-5C)(NIH HIV Reagent Program/BEI; ARP-3537), anti-FLAG (M2)(Sigma; F1804), anti-Myc (Sigma; C3956). The following secondary antibodies were used: goat anti-mouse IRDye 800CW (Li-COR; 926-32210), goat anti-mouse IRDye 680RD (Li-COR; 926-68070), goat anti-rabbit 800CW (Li-COR; 926-32211), goat anti-rabbit 680RD (Li-COR; 926-68071). Images were obtained with the Li-COR Odyssey CLx and analysis was performed with ImageStudio Lite software (Li-COR). PageRuler Prestained Protein Ladder, 10-180 kDa was used as protein standard (Thermo Fisher; 26616).

### Co-immunoprecipitation

Anti-HA antibody (Abcam; ab9110) or anti-FLAG antibody (M2)(Sigma; F1804) was added to 100 μg whole cell lysates, and the protein-antibody mixture was incubated for 1 hour at 4°C. 10 μL of Dynabeads Protein G (Invitrogen; 10007D) was added to the protein-antibody mixture for 1 hour at 4°C, and reaction mixtures were centrifuged at 1000 x g for 3 mins at 4°C. The supernatant was removed, and the pelleted beads were washed three times with cell lysis buffer. The bead fraction was resuspended with NuPAGE Reducing Agent (Invitrogen; NP0009) and Protein Loading Buffer (Li-COR; 928-40004), boiled for 5 mins at 95°C, and loaded into Criterion XT 12% polyacrylamide Bis-Tris gels (Bio-Rad; 3450117). SDS-PAGE and western blot analysis were performed as described above.

### Proximity Ligation Assay

In situ proximity ligation assay (PLA) was performed with the Duolink In Situ Red Starter Kit Mouse/Rabbit (Sigma; DUO92101) according to the manufacturer’s protocol. HEK293T cells were transfected and seeded (50,000 cells/well) in 8 well Mu-slide dish (Ibidi; 80826), fixed/permeabilized with Cytofix/Cytoperm Solution (BD; B554714) for 5 min and blocked with Duolink Blocking Solution (1X) for 1 hour at 37°C. Cells were then incubated with primary antibodies (rabbit anti-IFITM3 (Abcam; ab109429) and mouse anti-HA (Biolegend; 901533)) for 1 hour at room temperature. Cells were washed twice with Buffer A and subsequently incubated with the probes affinity purified Donkey anti-Rabbit IgG (anti-Rabbit PLUS, Sigma; DUO92002) and affinity purified Donkey anti-Mouse IgG (anti-Mouse PLUS, Sigma; DUO92004) for 1 hour at 37°C). After washing cells twice with Buffer A, oligonucleotide ligation was performed for 30 min at 37°C. Cells were washed two additional times with Buffer A, followed by incubation with amplification stock solution for 100 min at 37°C. After washing twice with Buffer B, Hoechst 33342 (Thermo Fisher; H3570) was added for 5 min at room temperature to label nuclei. Image acquisition was performed with a Leica Stellaris confocal fluorescence microscope.

### Confocal Immunofluorescence Microscopy

HEK293T were transfected and seeded (25,000 cells/well) in an 8 well Mu-slide dish (Ibidi; 80826), fixed/permeabilized with Cytofix/Cytoperm Solution (BD; 554722) for 5 min, and blocked with 3% bovine serum albumin. Cells were then incubated with primary antibodies (rabbit anti-IFITM3 (Abcam; ab109429) and mouse anti-HA (Biolegend; 901533)) for 1 hour at room temperature. After washing twice with Cyto Perm/Wash Solution (BD; 554723), cells were incubated with secondary antibodies (goat anti-mouse IgG (H+L) Secondary Antibody, Alexa Fluor 555 and goat anti-rabbit IgG (H+L) Secondary Antibody, Alexa Fluor 647). Nuclei were labeled with Hoechst 33342 and image acquisition was performed using a Leica Stellaris confocal microscope. Images were analyzed and processed using Fiji (Image J).

### Measurement of membrane fluidity by fluorescence lifetime imaging (FLIM)

HEK293T stably expressing IFITM3 or Empty Vector (previously described (32)) were untransfected or transfected with pBJ-97ZA Nef-HA (0.25 μg). Methyl-beta-cyclodextrin (MBCD) (Sigma; C4555) was added to HEK293T-IFITM3 cells at a final concentration of 5 mM for 1 hour at 37°C. Flipper-TR probe (Cytoskeleton; CY-SC020) was added to cells at a final concentration of 1 μM, in serum-free media, for 10 mins at 37°C. Media was replaced with Fluorobrite DMEM supplemented with 1% Pen-Strep and 10% FBS and imaging was performed on a Zeiss AG 880 laser confocal fluorescence microscope equipped with a live cell imaging environmental chamber and a Coherent Chameleon Vision II pulsed near infrared laser used for two-photon excitation, in conjunction with SPCM 64 FLIM acquisition software. Lifetimes were detected with an HPM-100-40 detector (Becker & Hickl) and collection time was set to 60 secs using SPC-QC-104 timing electronics (Becker & Hickl). Analysis was performed with SPCImage 8.8 software (Becker & Hickl). Raw images were uploaded and parameters were set as follows: spatial binning of 2 and threshold of 6; single exponential decay analysis. A region of interest tracing the periphery of each cell was established, and the lifetime of all pixels within the region of interest was averaged. Analyzed images were scaled, from red to blue, on a range of 1800 ps to 3800 ps.

### Statistical analysis

Tests for statistical significance were performed in GraphPad Prism.

## ACKNOWLEDGEMENTS

We would like to thank Chen Liang (McGill University) for providing the Tet ON SupT1-IFITM3 cells, Olivier Schwartz and Nicoletta Casartelli (Pasteur Institute) for providing full length pCH040, Richard Sloan (University of Edinburgh) for providing pNL4-3 based constructs encoding HIV-2 Nef and SIVmac239 Nef, Heinrich Gottlinger (University of Massachusetts) for providing pBJ-SF2 Nef-HA, pBJ-LAI Nef-HA, pBJ-97ZA Nef-HA, pBJ-90CF Nef-HA, pBJ-93BR Nef-HA, and pBJ-94UG Nef-HA constructs and pNL4-3 based viruses encoding Nef from primary isolates of HIV-1, Alan Rein (National Cancer Institute) for providing MLV plasmids and virus production assistance, John Guatelli (University of California San Diego) for providing pCl-Nef constructs encoding T/F Nef proteins, and Stephen Lockett (Frederick National Laboratory for Cancer Research) for providing technical support and training for fluorescence lifetime imaging.

