## Supplementary figures and images for "Restriction of HIV-1 infectivity by interferon and IFITM3 is counteracted by Nef"

### Supplemental Figure 1

Supplemental Figure 1

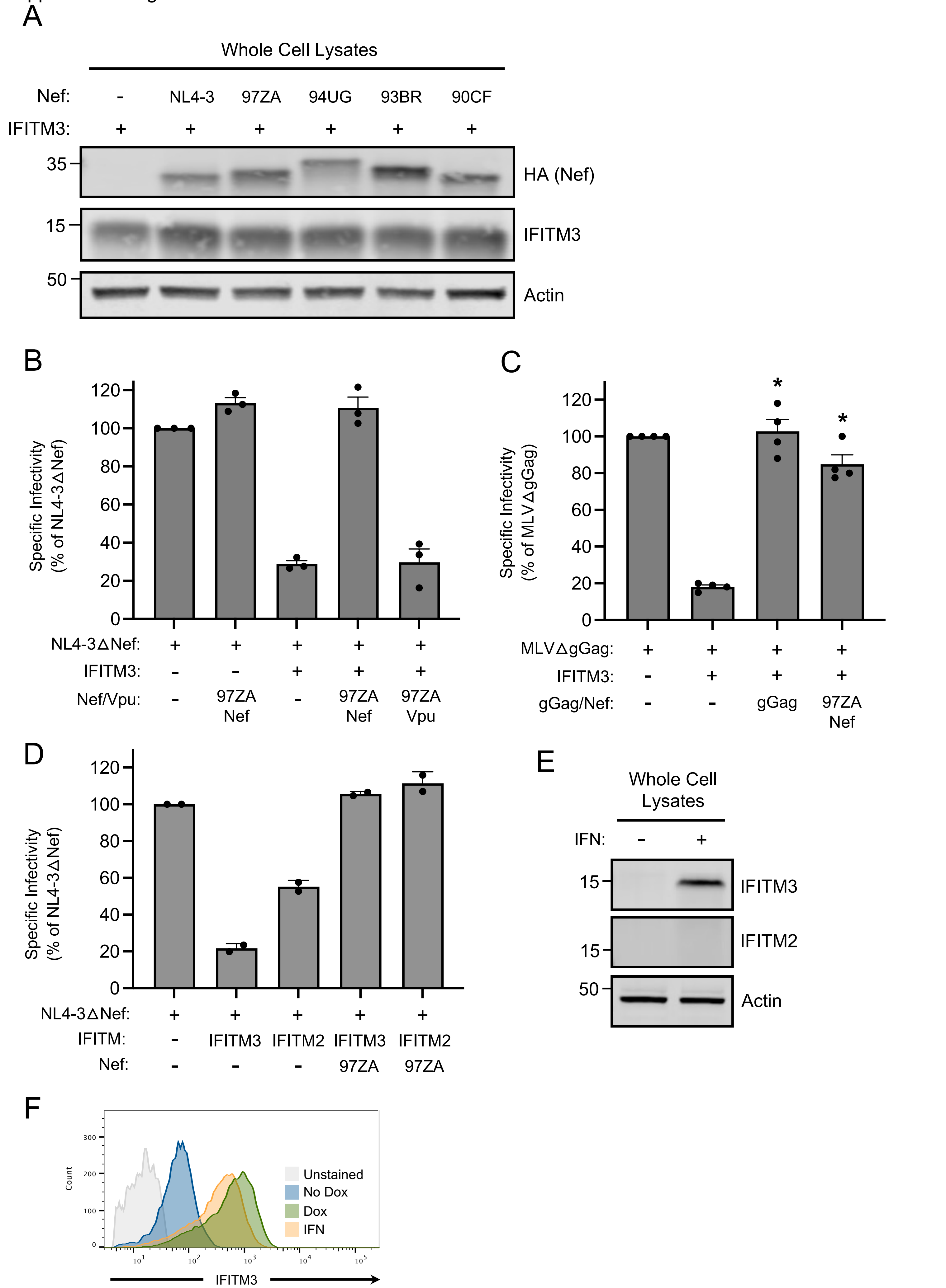

### Supplemental Figure 2

Supplemental Figure 2

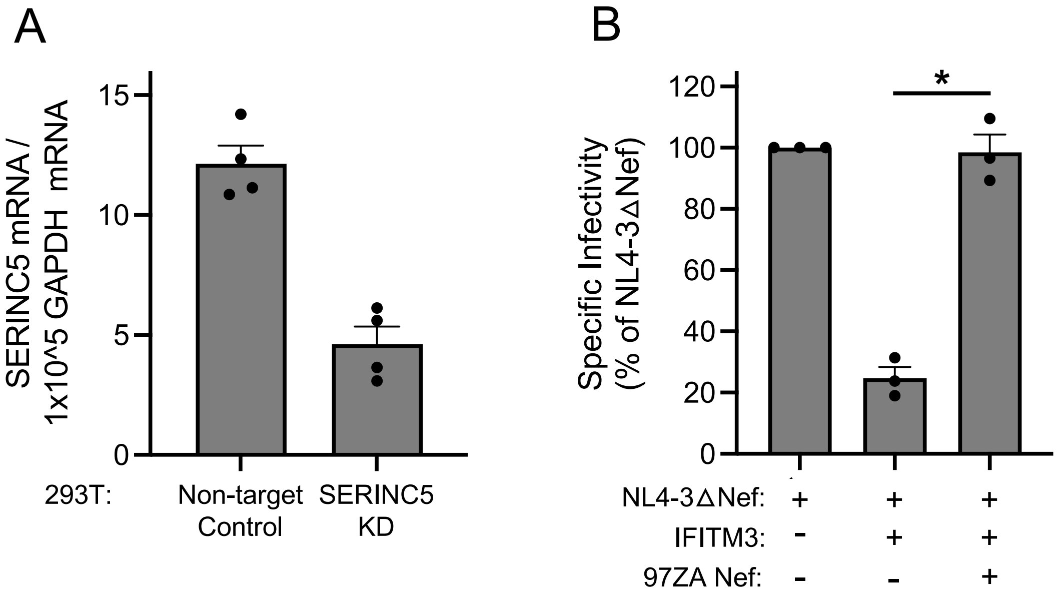

### Supplemental Figure 3

Supplemental Figure 3

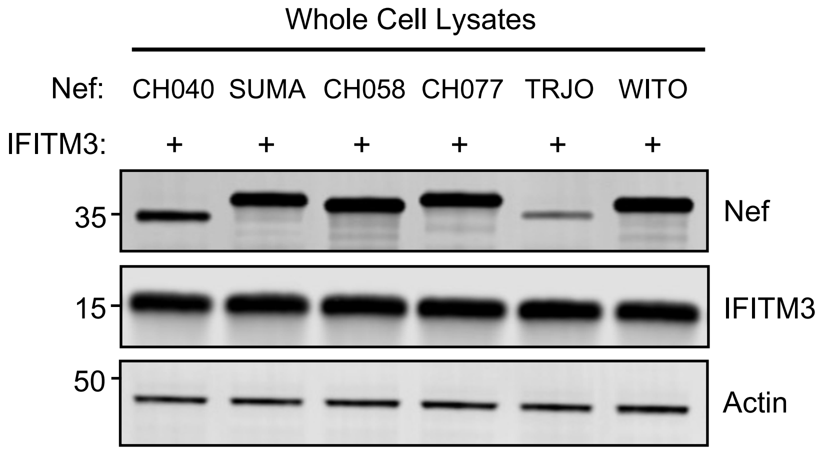
