## Supplemental File 1 for "Restriction of HIV-1 infectivity by interferon and IFITM3 is counteracted by Nef"

**Nef protein sequences from this study:**

>NL4-3
MGGKWSKSSVIGWPAVRERMRRAEPAADGVGAVSRDLEKHGAITSSNTAANNAACAWLEAQEEEEVGFPVTPQVPLRPMTYKAAVDLSHFLKEKGGLEGLIHSQRRQDILDLWIYHTQGYFPDWQNYTPGPGVRYPLTFGWCYKLVPVEPDKVEEANKGENTSLLHPVSLHGMDDPEREVLEWRFDSRLAFHHVARELHPEYFKNC

>SF2

MGGKWSKRSMGGWSAIRERMRRAEPRAEPAADGVGAVSRDLEKHGAITSSNTAATNADCAWLEAQEEEEVGFPVRPQVPLRPMTYKAALDISHFLKEKGGLEGLIWSQRRQEILDLWIYHTQGYFPDWQNYTPGPGIRYPLTFGWCFKLVPVEPEKVEEANEGENNSLLHPMSLHGMEDAEKEVLVWRFDSKLAFHHMARELHPEYYKDC

>LAI

MGGKWSKSSVVGWPTVRERMRRAEPAADGVGAASRDLEKHGAITSSNTAATNAACAWLEAQEEEEVGFPVTPQVPLRPMTYKAAVDLSHFLKEKGGLEGLIHSQRRQDILDLWIYHTQGYFPDWQNYTPGPGVRYPLTFGWCYKLVPVEPDKVEEANKGENTSLLHPVSLHGMDDPEREVLEWRFDSRLAFHHVARELHPEYFKNC

>97ZA012
MGGKWSKSSLVGWPNVRERMRRTEPAAEGVGAASRDLDKHGALTSSNTAHNNADCAWLQAQEETEEVGFPVRPQVPLRPMTYKAAIDLSFFLKEKGGLEGLIHSKRRQDILDLWVYHTQGYFPDWQNYTPGPGVRYPLTFGWCFKLVPVDPSEVEEANKGENNCLLHPMSQHGIEDAEREVLKWEFDSSLARRHIAREKHPEYYKDC

>NL43_97ZA012
MGGKWSKSSVIGWPAVRERMRRAEPAADGVGAVSRDLEKHGAITSSNTAANNAACAWLEAQEEEEVGFPVTPQVPLRPMTYKAAVDLSHFLKEKGGLEGLIHSQRRQDILDLWIYHTQGYFPDWQNYTPGPGVRYPLTFGWCFKLVPVDPSEVEEANKGENNCLLHPMSQHGIEDAEREVLKWEFDSSLARRHIAREKHPEYYKDC

>97ZA012_NL43
MGGKWSKSSLVGWPNVRERMRRTEPAAEGVGAASRDLDKHGALTSSNTAHNNADCAWLQAQEETEEVGFPVRPQVPLRPMTYKAAIDLSFFLKEKGGLEGLIHSKRRQDILDLWVYHTQGYFPDWQNYTPGPGVRYPLTFGWCYKLVPVEPDKVEEANKGENTSLLHPVSLHGMDDPEREVLEWRFDSRLAFHHVARELHPEYFKNC

>94UG114

MGGKWSKSSIVGWPAVRERMRRTEPAAEGVGAASRDLEKHGAITSSNTAQTNDACAWLEAQEEEEVGFPVRPQVPLRPMTYKEAVDLSHFLKEKGGLEGLVWSPKRQEILDLWVYHTQGFFPDWQNYTPGPGIRYPLTFGWCFELVPMEPKEVEENTEGEDNCLLHPINQHGMEDPEREVLVWRFNSRLAFEHKAKMKHPEYYKDC

>93BR020

MGGKWSKSSIVGWPAIRERMRRTPPTPPAAEGVGAVSQDLERRGAITSSNTRANNPDLAWLEAQEEDEVGFPVRPQVPLRPMTYKGAVDLSHFLKEKGGLEGLIYSKRRQEILDLWVYHTQGYFPDWQNYTPGPGIRYPLTMGWCFKLVPVDPEEVEKANEGENNCLLHPMSQHGMEDEDKEVLKWEFDSRLALRHIARERHPEYYQD

>90CF056

MGGKWSKSRMGGWSTIRERMRRAEPVAEGVGAVSRDLDRRGAVTINNTASTNRDAAWLEAQEDGEEVGFPVRPQVPLRPMTYKGAFDLSHFLKEKGGLDGLIYSKQRQDILDLWVYNTQGYFPDWQNYTPGPGERFPLTFGWCFKLVPVNPQEVEQANEGENNSLLHPMSLHGMEDDGREVLMWKFDSRLALTHLARVKHPEYKDC

>Acute_cladeC_Consensus
MGGKWSKSSIVGWPAVRERIRRTEPAAEGVGAASQDLDKHGALTSSNTAHNNADCAWLQAQEEEEEVGFPVRPQVPLRPMTYKAAFDLSFFLKEKGGLDGLIYSKKRQEILDLWVYHTQGFFPDWQNYTPGPGVRYPLTFGWCFKLVPVDPREVEEANKGENNCLLHPMSQHGMEDEEREVLKWKFDSSLARRHLARELHPEYYKDC

>HIV-2_BEN

MGASGSKKLSKHSRGLRERLLRARGDGYGKQRDASGGEYSQFQEESGREQNSPSCEGQQYQQGEYMNSPWRNPATERQKDLYRQQNMDDVDSDDDDLIGVPVTPRVPRREMTYKLAIDMSHFIKEKGGLQGMFYSRRRHRILDIYLEKEEGIIPDWQNYTHGPGVRYPMYFGWLWKLVSVELSQEAEEDEANCLVHPAQTSRHDDEHGETLVWQFDSMLAYNYKAFTLYPEEFGHKSGLPEKEWKAKLKARGIPYSE

>SIVmac239

MGGAISMRRSRPSGDLRQRLLRARGETYGRLLGEVEDGYSQSPGGLDKGLSSLSCEGQKYNQGQYMNTPWRNPAEEREKLAYRKQNMDDIDEEDDDLVGVSVRPKVPLRTMSYKLAIDMSHFIKEKGGLEGIYYSARRHRILDIYLEKEEGIIPDWQDYTSGPGIRYPKTFGWLWKLVPVNVSDEAQEDEEHYLMHPAQTSQWDDPWGEVLAWKFDPTLAYTYEAYVRYPEEFGSKSGLSEEEVRRRLTARGLLNMADKKETR

>CH040

MGGKWSKCSVVGWPSVRERMRRAEPAAEGVGAVSRDLEKHGAITSSNTAATNADCAWLEAQEEGEVGFPVRPQVPLRPMTFKGALDLSHFLKEKGGLEGLIYSQKRQDILDLWVYHTQGYFPDWQNYTPGPGTRFPLTFGWCFKLVPVDPGKVEEANKGENNCLLHPMSQHGMDDPEREVLVWRFDSSLAFRHVARELHPEYYKNC

>CH058

MGGKWSKRSVPGWADVRERMRRTEPRTEPAADGVGAVSRDLEKHGAITSSNTAANNPDCAWLEAQEEEEEVGFPVRPQVPLRPMTYKGALDLSHFLKEKGGLEGLIHSQKRQDILDLWVYHTQGYFPDWQNYTPGPGTRYPLTFGWCFKLVPVDPEKVEEANTGENISLLHPMSQHGMDDPEKEVLKWTFDSHLAFHHMARELYPEYYKN

>CH077

MGGKWSKFAGWPAVRERMRRAGARERRRRDEPAAVGVGPASQDLAKHGAITSSNTVSNNADCAWLEAQEEEEEVGFPVRPQVPVRPMTYKAALDLSHFLKEKGGLEGLIYSQQRKDILDLWVYNTQGFFPDWQNYTPGPGPRFPLTFGWCFKLVPVEPEEVEKANEGENNCLLHPMSQHGTDDPEKEVLAWRFDSRLAFQHVAREIHPEFYKDC

>SUMA

MGGKWSKSRGVGWSTIREKMRRAEPAAEPAAEGVGAVSRDLEKHGAITNSNTAATNADVAWLEAQEDEEVGFPVRPQVPLRPMTYKGAFDLSHFLKEKGGLEGLIYSRKRQEILDLWVYHTQGYFPDWQNYTPGPGIRYPLTFGWCFKLVPVEPEEVEKANEGESNCLLHPMSQHGMDDPEKEVLVWKFDSRLAFHHMARELHPEYYKDC

>TRJO

MGGKWSKRSVVGWPKVRERMRRVEPAADGVGAVSRDLDQRGAVTINNTPANNDTCAWLEAQEDEDVGFPVRPQVPLRPMTFKGALDLSHFLKEQGGLDGLIYSQKRQEILDLWIYHTQGYFPDWGNYTPGPGIRYPLTFGWCFKLVPVDPDEVEKANEGENNCLLHPMSQHGMDDPEKEVLMWKFDSMLAFQHKARELYPDYYKDC

>WITO

MGGKWSKSWKIGWPTVRERMRRAEPEPAAVGVGAVSRDLERHGAVTSSNTATNNADSAWLEAQAQEEDNEVGFPVRPQVPVRPMTYKAAVDLSHFLKEKGGLDGLIYSQQRQDILDLWVYNTQGFFPDWQNYTPGPGTRYPLTFGWCYKLVPVEPEEVEKANEGENNSLLHPMGLHGMDDPEKEVLMWKFDSRLAFHHMAREKHPEFYKDC

| **Nef** | **Accession** |
| --- | --- |
| NL4-3 | AGL78171.1 |
| SF2 | P03407.3 |
| LAI | P03406.3 |
| 97ZA012 | AAK30998.1 |
| 94UG114 | AAC97574.1 |
| 93BR020 | AAA99884.1 |
| 90CF056 | AF005496.1 |
| HIV-2 Ben | M30502.1 |
| SIVmac239 | AAB99967.1 |
| CH040 | ACR51130.1 |
| CH058 | QZK27722.1 |
| CH077 | QZK27743.1 |
| SUMA | ACR52828.1 |
| TRJO | AAG34603.1 |
| WITO | ACR52994.1 |
